# Renewable Fatty Acid Ester Production in *Clostridium*

**DOI:** 10.1101/2020.03.29.014746

**Authors:** Jun Feng, Jie Zhang, Yiming Feng, Pixiang Wang, Pablo Jiménez-Bonilla, Yanyan Gu, Junping Zhou, Zhong-Tian Zhang, Mingfeng Cao, Zengyi Shao, Ilya Borovok, Haibo Huang, Yi Wang

**Affiliations:** Department of Biosystems Engineering, Auburn University, Auburn, Alabama 36849, USA; Center for Bioenergy and Bioproducts, Auburn University, Auburn, AL 36849, USA; Department of Food Science and Technology, Virginia Tech, Blacksburg, VA 24061, USA; School of Chemistry, National University (UNA), Heredia, Costa Rica; Department of Chemical and Biological Engineering, Iowa State University, Ames, IA, 50011, USA; NSF Engineering Research Center for Biorenewable Chemicals, Iowa State University, Ames, IA, 50011, USA; The School of Molecular Cell Biology and Biotechnology, Faculty of Life Sciences, Tel Aviv University, Ramat Aviv, 6997801 Tel Aviv, Israel

**Author notes:** Corresponding author: Haibo Huang, Department of Food Science and Technology, Virginia Tech, 1230 Washington St. SW, Blacksburg, VA 24061, USA, Yi Wang, Department of Biosystems Engineering, Auburn University, 350 Mell Street, Auburn, AL, 36849 USA. These authors contributed equally to this work (Jun Feng, Jie Zhang).

**Keywords:** clostridia, biofuels and biochemicals, fatty acid ester, butyl acetate, butyl butyrate, CRISPR-Cas9, prophages

## Abstract

Production of renewable chemicals through biological routes is considered as an urgent solution for fossil energy crisis. However, endproduct toxicity inhibits microbial performance and is a key bottleneck for biochemical production. To address this challenge, here we report an example of biosynthesis of high-value and easy-recoverable derivatives to alleviate endproduct toxicity and enhance bioproduction efficiency. By leveraging the natural pathways in solventogenic clostridia for co-producing acyl-CoAs, acids and alcohols as precursors, through rational screening for host strains and enzymes, systematic metabolic engineering— including rational organization of ester-synthesizing enzymes inside of the cell, and elimination of putative prophages, we developed strains that can produce 20.3 g/L butyl acetate and 1.6 g/L butyl butyrate respectively, which were both the unprecedented levels in microbial hosts. Techno-economic analysis indicated a production cost of $986 per metric tonne for butyl acetate production from corn stover comparing to the market price of $1,200-1,400 per metric tonne of butyl acetate, suggesting the economic competitiveness of our developed bioprocess. Our principles of selecting the most appropriate host for specific bioproduction and engineering microbial chassis to produce high-value and easy-separable endproducts are highly applicable to other bioprocesses, and could lead to breakthroughs in biofuel/biochemical production and general bioeconomy.

## Main

Although tremendous efforts have been invested for biofuel and biochemical research, it is still challenging to generate robust microbial strains that can produce target products at desirable levels^1^. One key bottleneck is the intrinsic toxicity of endproducts to host cells^2^. Therefore, the production of high-value bioproducts which can be easily recovered from fermentation might be a solution to tackle the bottleneck in bioproduction. Fatty acid esters, or mono-alkyl esters, can be used as valuable fuels such as diesel components or specialty chemicals for food flavoring, cosmetic and pharmaceutical industries^3^. It is projected that the US market demand for fatty acid esters could reach $4.99 billion by 2025^4^. In addition, esters, with fatty acid and alcohol moieties, are generally hydrophobic and can easily separate from fermentation; thus the production of ester can help mitigate endproduct toxicity to host cells and efficient bioproduction can be achieved.

Conventionally, esters are produced through Fischer esterification which involves high temperature and inorganic catalysts^5,6^. The reaction consumes a large amount of energy and generates tremendous wastes, and thus is not environmentally friendly^5^. On the other hand, ester production through biological routes is becoming more and more attractive because it is renewable and environmentally benign. There are two primary biological pathways for ester production: one is through esterification of fatty acid and alcohol catalyzed by lipases^7^, and the other is based on condensation of acyl-CoA and alcohol catalyzed by alcohol acyl transferases (AATs)^5^. Previously, lipases from bacteria or fungi have been employed for catalyzing esterification for ester production^8^. For instance, lipase from *Candida sp*. has been recruited to drive production of butyl butyrate (BB) with *Clostridium tyrobutyricum. C. tyrobutyricum*, a natural hyper-butyrate producer, could generate 34.7 g/L BB with supplementation of lipase and butanol^9^. However, in such a process, the supplemented enzyme accounts a big cost and meanwhile the operation needs to be carefully managed to achieve the optimum performance of esterification^9^. Therefore, a whole microbial cell factory able of ester production in one pot is highly desired. *Saccharomyces cerevisiae* has been reported to produce various esters with its native AATs, but generally at very low levels (< 1 g/L)^10^. Rodriguez et al. metabolically engineered *E. coli* to produce esters by introducing heterologous AATs^5^. Although the production of some acetate esters can reach decent levels (such as 17.2 g/L isobutyl acetate), the production of most of the esters was rather low, probably due to the unavailability of intrinsic substrates/precursors and limited tolerance of *E. coli* to organic endproducts.

In this study, we report highly efficient fatty acid ester production to unprecedented levels using engineered clostridia. We selected solventogenic clostridia to take advantage of their natural pathways for co-producing acyl-CoAs (acetyl-CoA and butyryl-CoA), fatty acids (acetate and butyrate), and alcohols (ethanol and butanol), either as intermediates or endproducts; we hypothesized that clostridia can be excellent microbial platforms to be engineered for efficient ester production by introducing heterologous AATs and/or lipase genes. Indeed, through rational screening for host strains (from four well-known clostridial species) and enzymes (alcohol acyl transferases and lipase), systematic metabolic engineering— including rational organization of ester-synthesizing enzymes inside of the cell, and elimination of putative prophages, we ultimately obtained two strains which can produce 20.3 g/L butyl acetate (BA) and 1.6 g/L BB respectively in extractive batch fermentations. These production levels were both highest in record.

## Results

### Screening of host strains and genes for ester production

We considered clostridia as ideal platforms for ester production thanks to their intrinsic intermediates (fatty acids, acyl-CoAs, and alcohols) serving as precursors for ester biosynthesis (Fig. 1). We hypothesized that different flux levels of these precursors within various clostridial strains would make a big difference for the specific type(s) of ester production. Therefore, we selected five strains (from four representative species) including *C. tyrobutyricum* Δ*cat1::adhE1*^11^, *C. tyrobutyricum* Δ*cat1::adhE2*^11^, *C. pasteurianum* SD-1^12^, *C. saccharoperbutylacetonicum* N1-4-C^13^ and *C. beijerinckii* NCIMB 8052^14^ to evaluate their capabilities for ester production through metabolic engineering (Table S1). We included both *C. tyrobutyricum* Δ*cat1::adhE1* and Δ*cat1::adhE2* here because they produce different levels (and thus ratios) of butanol and ethanol^11,15^. Esters can be synthesized either through esterification of acid and alcohol catalyzed by lipase, or through condensation of acyl-CoA and alcohol catalyzed by AATs (Fig. 1). Previously, lipase B (CALB) from *Candida antarctica* has been employed for efficient ester production through esterification^9,16^. In addition, four AATs including VAAT^*17,18*^, SAAT^17^, ATF1^3,10^, EHT1^19^ have been recruited for ester production in various hosts^5,20-22^. Therefore, here, we evaluated all these genes in our clostridial hosts for ester production.

**Fig. 1.**
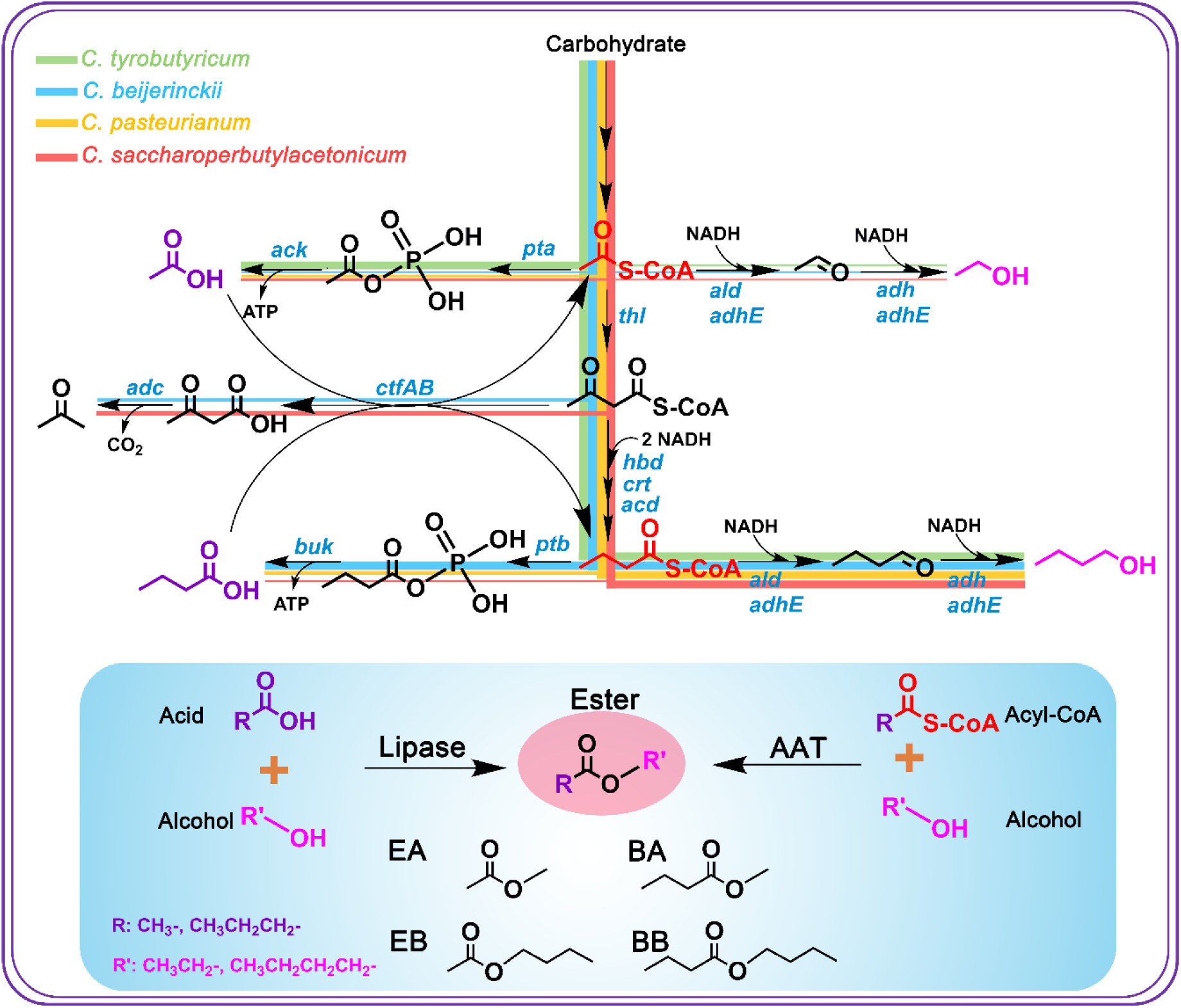
Engineering of solventogenic clostridia for fatty acid ester production. **Top:** Five strains out of four representative clostridial species were selected and evaluated as the host to be engineered for ester production in this study. We hypothesized that the different metabolic fluxes within different strains would make a big difference for the desirable ester production. The metabolic pathways of the four different species were represented in four different colors. **Bottom:** Fatty acid esters could be synthesized through two primary biological pathways: one is through the esterification of fatty acid and alcohol catalyzed by lipases, and the other is through the condensation of acyl-CoA and alcohol catalyzed by alcohol acyl transferases (AATs). Key genes in the pathway: *pta*, phosphotransacetylase; *ack*, acetate kinase; *thl*, thiolase; *hbd*, beta-hydroxybutyryl-CoA dehydrogenase; *crt*, crotonase; *bcd*, butyryl-CoA dehydrogenase; *adh*, alcohol dehydrogenase; *adhE*, Aldehyde-alcohol dehydrogenase; *adc*, acetoacetate decarboxylase; *ctfAB*, CoA transferase; *ptb*, phosphotransbutyrylase; *buk*, butyrate kinase; *ald*, aldehyde dehydrogenase.

Six plasmids (pMTL-P_*cat*_-*vaat*, pMTL-P_*cat*_-*saat*, pMTL-P_*cat*_-*atf1*, pMTL-P_*cat*_-*eht1*, pMTL-P_*cat*_-*lipaseB* as well as pMTL82151 as the control) were individually transformed into *C. saccharoperbutylacetonicum* N1-4-C, *C. pasteurianum* SD-1, *C. tyrobutyricum cat1::adhE1* and *cat1::adhE2* respectively. While pTJ1-P_*cat*_-*vaat*, pTJ1-P_*cat*_-*saat*, pTJ1-P_*cat*_-*atf1*, pTJ1-P_*cat*_-*ehtl* and pTJ1-P_*cat*_-*lipaseB* as well as pTJ1 were transformed into *C. beijerinckii* 8052. Fermentations were performed (Fig. 2a), and results were shown in Fig. 2b. Four types of esters were detected: EA, BA, ethyl butyrate (EB) and BB. Interestingly, control strains with the empty plasmid (pMTL82151 or pTJ1) also produced noticeable EA, BA and BB. This could be because: 1) the endogenous lipase in clostridia can catalyze ester production^13^; 2) *catP* on pMTL82151 encoding a chloramphenicol acetyltransferase (belonging to the same class of enzymes as AATs) has AAT activities^5,23^.

**Fig. 2.**
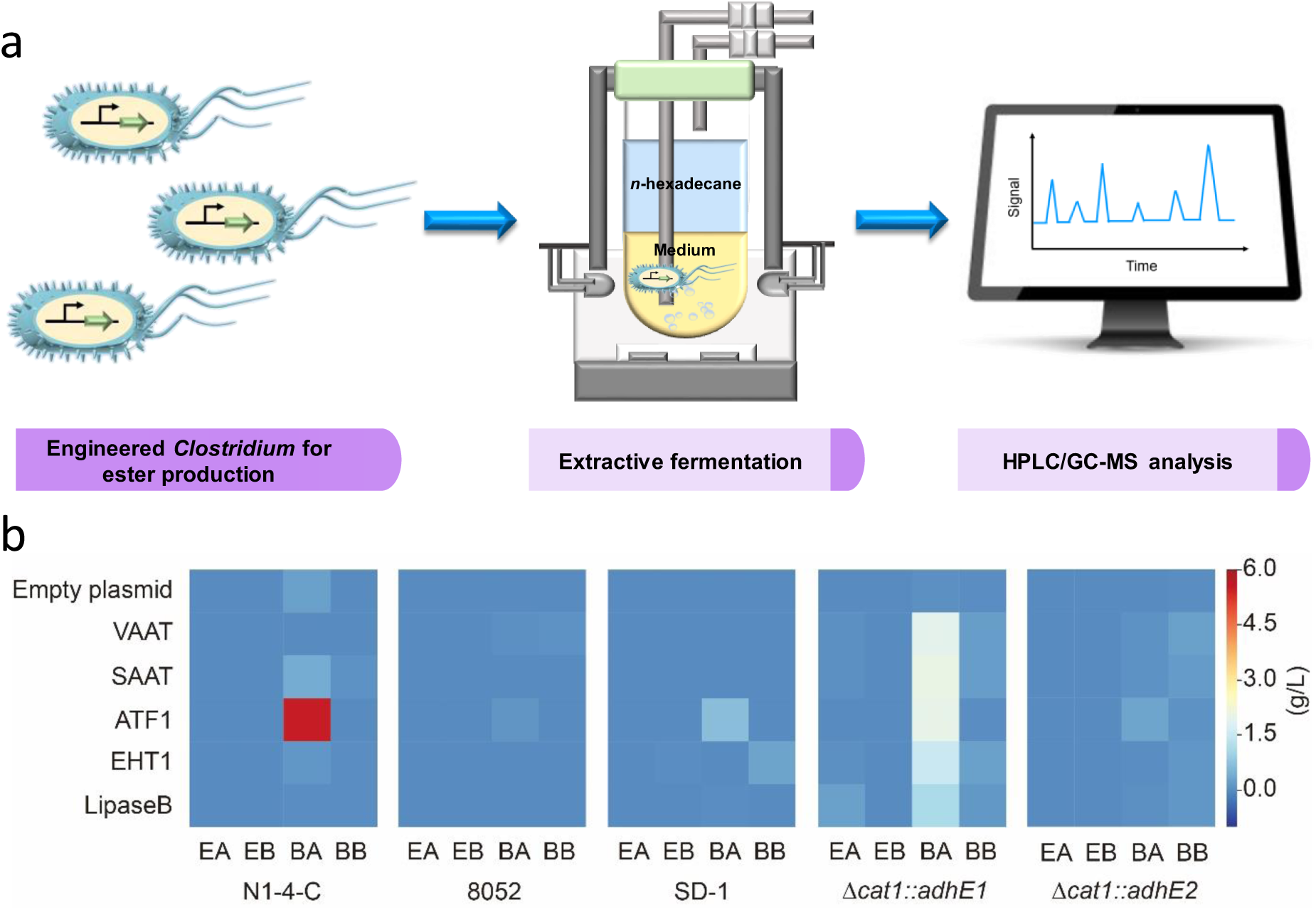
Screening of strains and enzymes for ester synthesis. (a) Schematic representation of devices and procedures for ester fermentation with serum bottle. (b) Heatmap results showed the ester production in various *Clostridium* strains with the overexpression of different enzymes. Empty plasmid: overexpression of pMTL82151 (for *C. saccharoperbutylacetonicum* N1-4-C, *C. pasteurianum* SD-1, *C. tyrobutyricum cat1::adhE1* and *cat1::adhE2*) or pTJ1 (for *C. beijerinckii* 8052) as the control; EA: ethyl acetate; EB: ethyl butyrate; BA: butyl acetate; BB: butyl butyrate. Scale bar on the right is in g/L.

Based on the results, it could be concluded that ATF1 is more favorable for BA production. All strains with *atf1* produced higher levels of BA compared to the same strain but with the overexpression of other genes (Fig. 2b). While VAAT, SAAT and EHT1 seemed to have better activities for BB production. Among all the strains, *C. saccharoperbutylacetonicum* FJ-004 produced the highest titer of 5.5 g/L BA. This is the highest BA production that has ever been reported through microbial fermentation. *C. acetobutylicum* CaSAAT (with the overexpression of *saat* from *Fragaria xananassa*) was reported to produce 8.37 mg/L BA^22^. While this manuscript was prepared, a newly engineered *C. diolis* strain was reported to produce 1.37 g/L BA^24^. The highest BB production of 0.3 g/L was observed in *C. pasteurianum* J-5 with the overexpression of *eht1*. This is also the highest BB production reported so far directly from glucose with engineered microorganisms, which is significantly higher than the recently reported 50.07 mg/L in an engineered *C. acetobutylicum*^22^. *Couple of our engineered strains could also produce small amount of EB with the highest of 0*.*02 g/L been observed in C. pasteurianum* J-5. With the *lipaseB* overexpression, *C. tyrobutyricum* JZ-6 could generate 0.3 g/L EA. This was significantly higher than other strains tested in this work (mostly < 0.01 g/L). As we reported previously, the mother strain *C. tyrobutyricum* Δ*cat1::adhE1* could produce 20.8 g/L acetate and 5.3 g/L ethanol (precursors for EA synthesis) during a batch fermentation, which might have enabled high-level EA production in *C. tyrobutyricum* JZ-6^11^.

The production levels of BA and BB achieved above are both significantly higher than the previously reported in microbial hosts. In comparison, the BA level is much higher than BB level, and thus has greater potentials towards economic viability. Therefore, in the following steps, we primarily focused on systematic metabolic engineering of the strain for further enhanced BA production.

### Deletion of *nuoG* increased BA production

Enhancement of the pool of precursors is one common strategy to improve the production of targeted bioproduct. Butanol and acetyl-CoA are the two precursors for BA production. Therefore, we set out to increase butanol synthesis to improve BA production. The *nuoG* gene encodes the NADH-quinone oxidoreductase subunit G, which is a subunit of the electron transport chain complex I^25^. NADH-quinone oxidoreductase can oxidize NADH to NAD^+^ and transfer protons from cytoplasm to periplasm to form a proton gradient between periplasm and cytoplasm, which can then contribute to the energy conversion^25^ (Fig. S1). It has been reported that inactivation of *nuoG* could increase both glucose consumption and butanol production in *C. beijerinckii*^26^. *In this study, we hypothesized that by deleting nuoG*, BA production would be boosted because of the potentially increased butanol production. Thus, we deleted *nuoG* (*Cspa_c47560*) in N1-4-C and generated FJ-100. Further, FJ-101 was constructed based on FJ-100 for BA production. Results demonstrated that, although butanol production in FJ-100 was only slightly improved (16.5 g/L vs. 15.8 g/L in N1-4-C; Fig. S2), BA production in FJ-101 was remarkably enhanced compared to FJ-004 (7.8 g/L vs. 5.5 g/L). Based on our experiences, because *C. saccharoperbutylacetonicum* N1-4 (or N1-4-C) mother strain can naturally produce very high level butanol, it was generally very difficult to further improve butanol production in *C. saccharoperbutylacetonicum* through simple metabolic engineering strategies^27^. This is likely the case in FJ-100 (comparing to N1-4-C). However, the increased NADH availability with *nuoG* deletion in FJ-101 would enable an enhanced ‘instant’ flux/availability of butanol which would serve as a precursor for BA production and thus enhance BA production in FJ-101. In this sense, the total butanol generated during the process (including the fraction serving as the precursor for BA production and the other fraction as the endproduct) in FJ-101 would be actually much higher than in FJ-004.

### Enhancement of acetyl-CoA availability to improve BA production

At the end of fermentation with FJ-101, there was still 7.6 g/L of butanol remained. This suggested that limited availability of intracellular acetyl-CoA was likely the bottleneck for the further improvement of BA production. To enhance the availability of acetyl-CoA, we firstly introduced isopropanol synthesis (from acetone) pathway (Fig. 3a). We hypothesized that this could pull flux from acetone to isopropanol, and thus boost the transferring of CoA from acetoacetyl-CoA to acetate driven by the CoA transferase, resulting in increased instant availability of acetyl-CoA. The *sadh* gene in *C. beijerinckii* B593 encoding a secondary alcohol dehydrogenase can convert acetone into isopropanol^28^. The *hydG* gene in the same operon as *sadh* encodes a putative electron transfer protein. It has been demonstrated to play important roles for the conversion of acetone into isopropanol^29,30^. In this study, either *sadh* alone or the *sadh*-*hydG* gene cluster was integrated into the chromosome of FJ-100, generating FJ-200 and FJ-300, respectively. Further, by introducing pMTL-cat-*atf1*, FJ-201 and FJ-301 were obtained. Compared to FJ-101, about 50% of the acetone could be converted into isopropanol in FJ-201, while ∼95% of the acetone in FJ-301 could be converted into isopropanol. The total titers of acetone plus isopropanol in FJ-301 and FJ-201 were 6.6 and 5.1 g/L respectively, both of which were higher than 4.8 g/L acetone in FJ-101. More significantly, BA production in FJ-201 and FJ-301 has been remarkably increased compared to FJ-101, and reached 10.1 and 12.9 g/L, respectively, likely due to enhanced ‘regeneration’ of acetyl-CoA as described above (Fig. 3b).

**Fig. 3.**
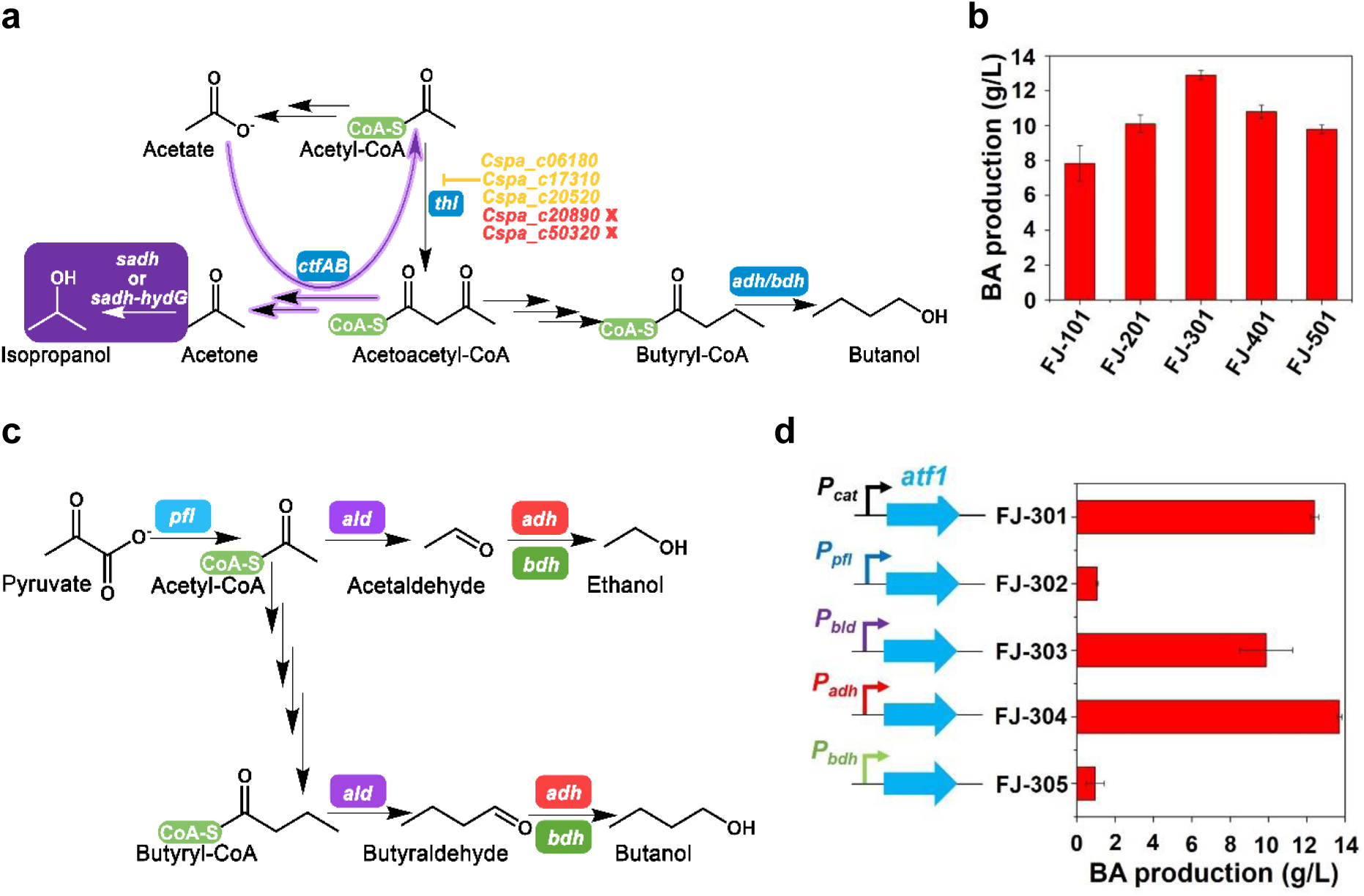
Enhancing BA production through increasing the availability of acetyl-CoA (a & b) and dynamically expressing the *atf1* gene (c & d). (a) Introduction of isopropanol synthesis pathway (shaded in purple) and deletion of thiolase genes. The pathways in purple arrows represent the ‘regeneration’ of actyl-CoA. There are five annotated genes encoding thiolase in *C. saccharoperbutylacetonicum*, only two (in red) of which could be deleted (see Supplementary materials). (b) The fermentation results for BA production with various mutant strains corresponding to the genetic manipulations in (a). The reported value is mean ± SD. (c) Four promoters associated with the biosynthesis of acetyl-CoA or alcohols in the pathway were selected to drive the expression of *atf1*. (d) The fermentation results for BA production with various mutant strains in which different promoters were used to drive the expression of *atf1* as illustrated in (c). The reported value is mean ± SD. BA: butyl acetate. Key genes in the pathway: *adh*, alcohol dehydrogenase; *bdh*, butanol dehydrogenase; *ctfAB*, CoA transferase; *sadh*, secondary alcohol dehydrogenase; *hydG*, putative electron transfer protein; *pfl*, pyruvate formate lyase; *ald*, aldehyde dehydrogenase.

### Dynamic expression of *atf1* enhanced BA production

In our engineered strain, BA is synthesized through condensation of butanol and acetyl-CoA catalyzed by ATF1. The constitutively high expression of ATF1would not necessarily lead to high BA production. For example, BA production in FJ-008 in which *atf1* was driven by the constitutive strong promoter P_*thl*_ from *C. tyrobutyricum* was actually much lower (3.5 g/L vs 5.5 g/L) than in FJ-004 in which *atf1* was expressed under the promoter P_*cat*_ from *C. tyrobutyricum*. We hypothesized that, in order to obtain more efficient BA production, the synthesis of ATF1 should be dynamically controlled and thus synchronous with the synthesis of precursors (butanol or acetyl-CoA). Therefore, for the next step, we attempted to evaluate various native promoters for *atf1* expression, and identify the one(s) that can enable an appropriately dynamic expression of ATF1 and lead to enhanced BA production.

Four promoters associated with the synthesis of BA precursors were selected to drive the *atf1* expression, and four strains were constructed correspondingly for BA production (Fig. 3c). Fermentation results were shown in Fig. 3d. Indeed, distinct results for BA production were observed in these strains. The BA level was only 1.0 and 1.1 g/L in FJ-305 and FJ-302 with P_*bdh*_ and P_*pfl*_ for *atf1* expression, respectively. P_*ald*_ is an important promoter in *C. saccharoperbutylacetonicum*, which can sense the acidic state and switch cell metabolism from acidogenesis to solventogenesis^31^. FJ-303 with the *atf1* expression driven by P_*ald*_ produced 9.9 g/L BA, which was still 20% lower than in FJ-301. Interestingly, FJ-304 in which *atf1* was expressed under P_*adh*_ produced 13.7 g/L BA, which was about 10.5% higher than in FJ-301. Based on the results, we speculated that the promoter of *adh* (which encodes the key enzyme catalyzing butanol and ethanol production) might have resulted in the more appropriate dynamic expression of *atf1* in line with the flux of the precursors and thus led to enhanced BA production. On the other hand, ethanol production in FJ-304 was also lower than in FJ-301 (0.3 g/L vs 0.6 g/L). The enhanced BA production in FJ-304 consumed more acetyl-CoA, and thus decreased production of ethanol, which also needed acetyl-CoA as the precursor.

### Rational organization of BA-synthesis enzymes to enhance BA production

Rational organization of enzymes associated with the synthesis of target product is an effective strategy to improve the bioproduction^32^. In this study, we evaluated several such approaches to increase BA production (Fig. 4).

**Fig. 4.**
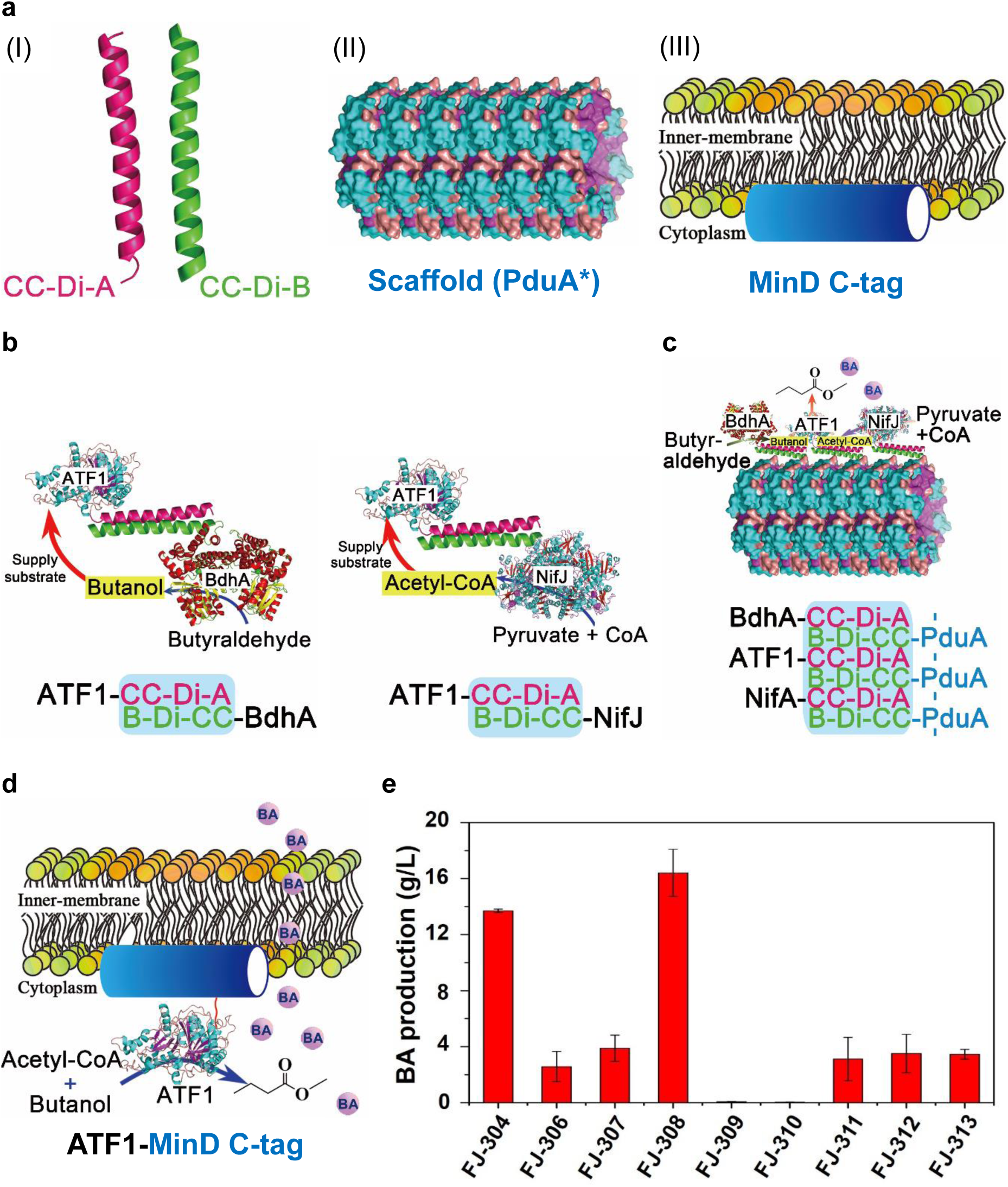
Enhancing BA production through rational organization of the enzymes associated with BA synthesis. (a) Biological components evaluated in this study for rational organization of enzymes: (I) CC-Di-A and CC-Di-B tags, (II) PduA* scaffold, (III) MinD C-tag. (b) Schematic representation of assembling two of the three enzymes for BA synthesis with the CC-Di-A and CC-Di-B tags. (c) Schematic representation of organizing the three enzymes for BA synthesis onto the PduA* formed scaffold. (d) Schematic representation of using the MinD C-tag to draw ATF1 onto the cell membrane. (e) The fermentation results for BA production with various mutant strains corresponding to the genetic manipulations from (b)-(d). The reported value is mean ± SD. BA: butyl acetate.

PduA* protein, derived from Citrobacter freundii Pdu bacterial microcompartment could form filaments in bacteria like E. coli^32^. The CC-Di-A and CC-Di-B are designed parallel heterodimeric coiled coils and two proteins with each of these self-assembling tags could combine and shorten the catalytic distance. The enzymes (from the same metabolic pathway), tagged with one of the coiled coils (CC-Di-A or CC-Di-B) would attach onto the formed intracellular filaments (its PduA* was tagged by the other coiled coil); thus, the organized enzymes on the filaments would improve the catalytic efficiency of the target metabolic pathway. MinD is a membrane-associated protein and the localization of MinD is mediated by an 8-12 residue C-terminal membrane-targeting sequence. The proteins with MinD C-terminal sequence were able to be drawn to the cell membrane^32, 33^. Thus, the application of MinD C-tag can facilitate the secretion of target product and enhance its production by mitigating the intracellular toxicity as well as promoting the catalyzing process (Fig. 4a).

To evaluate whether the organization of enzymes could improve BA production in our strain, three strategies were recruited: 1) assembling two of the three enzymes (enzymes associated with BA synthesis: NifJ (related to acetyl-CoA synthesis), BdhA (related to butanol synthesis) and ATF1) with the CC-Di-A and CC-Di-B tags; 2) organizing the three enzymes onto PduA* formed scaffold; or 3) introducing MinD C-tag to draw ATF1 onto the cell membrane.

Firstly, we assembled enzymes for BA synthesis by adding the CC-Di-A tag to ATF1 and the CC-Di-B tag to NifJ and BdhA (Fig. 4b). Fermentation results showed that the addition of CC-Di-A tag to the C-terminus of ATF1 in FJ-306 had significantly negative effects on BA synthesis with only 2.6 g/L BA was produced (Fig. 4e). The assembly of ATF1 together with NifJ or BdhA had even severer negative effects on BA synthesis and BA production was only 0.06 and 0.03 g/L in FJ-309 and FJ-310, respectively. Further, we organized ATF1, NifJ and BdhA onto the PduA* scaffold. PduA* was tagged with CC-Di-B, while the other three enzymes were tagged with CC-Di-A (Fig. 4c). The generated FJ-311 (harboring the scaffold and ATF1-CC-Di-A) produced 3.1 g/L BA, while FJ-312 (harboring the scaffold, ATF1-CC-Di-A and NifJ-CC-Di-A) and FJ-313 (harboring the scaffold, ATF1-CC-Di-A, NifJ-CC-Di-A and BdhA-CC-Di-A) produced slightly higher amount of BA both at 3.5 g/L. The scaffold seemed to have some positive effects on BA synthesis but didn’t work as efficient as it was reported in other studies^32^. We speculate that the coiled coils tags (CC-Di-A or CC-Di-B) might severely impair the catalytic activity of ATF1. Furthermore, the assembly of ATF1 together with NifJ or BdhA could further inhibit the activity of ATF1, thus resulting in significant decrease in BA production in the corresponding strains.

Moreover, we evaluated the effect of the introduction of MinD C-tag (to draw ATF1 onto the cell membrane) on BA production (Fig. 4d). Fermentation results showed that the corresponding FJ-308 strain could produce 16.4 g/L BA, which was 20% higher than FJ-304 (Fig. 4e). The FJ-307 strain (with CC-Di-A-Atf1-MinD) could also produce higher BA of 3.9 g/L compared to FJ-306 (2.6 g/L). All these results indicated that the addition of the cell membrane associated motif to draw ATF1 onto the cell membrane could facilitate the BA excretion and mitigate the intracellular toxicity and therefore enhance BA production. However, the assembly of BA synthetic enzymes or the organization of relevant enzymes onto the scaffold would significantly decrease BA production.

### Elimination of prophages increased BA (and BB) production

During our fermentations, we noticed that the ester production of the strains was not stable and could be varied from batch to batch. Our industrial collaborator also observed that *C. saccharoperbutylacetonicum* often had instable performance for butanol production in continuous fermentations (data not shown). It has been reported that the N1-4 (HMT) strain contains a temperate phage named HM T which could release from the chromosome even without induction^34^. In addition, the N1-4 (HMT) strain can produce a phage-like particle clostocin O upon the induction with mitomycin C^35^. We hypothesized that the instability of fermentations with *C. saccharoperbutylacetonicum* might be related to the prophages, and the deletion of prophages would enable more stable and enhanced production of desired endproducts. We identified five prophage-like genomes (referred here as P1-P5 respectively) within the chromosome of N1-4 (HMT) (Fig. 5a)^36^. Based on systematic evaluation through individual and combinatory deletion of the prophages, we demonstrated that P5 is responsible for the clostocin O synthesis (Figs. 5f, h & S5)^35^, and further confirmed that P1 was the HM T phage genome (Figs. 5g & S7). However, the phage image was different from what was described before^37^. It was more like HM 7 (a head with a long tail), rather than HM 1 (a head with multiple short tails). This is the first time that an image of the HM T phage has been reported. Both ΔP1234 (with the deletion of P1-P4) and ΔP12345 (with the deletion of P1-P5) exhibited improved cell growth and enhanced butanol production (Figs. 5b-5e, Figs. S3, S8 & S9). ΔP12345 should be a more stable platform as there is no cell lysis at any induction conditions with mitomycin C or norfloxacin (Fig. 5i). While ΔP1234 showed similar growth and even slightly higher butanol production compared to ΔP12345 (Figs. 5d & 5e).

**Fig. 5.**
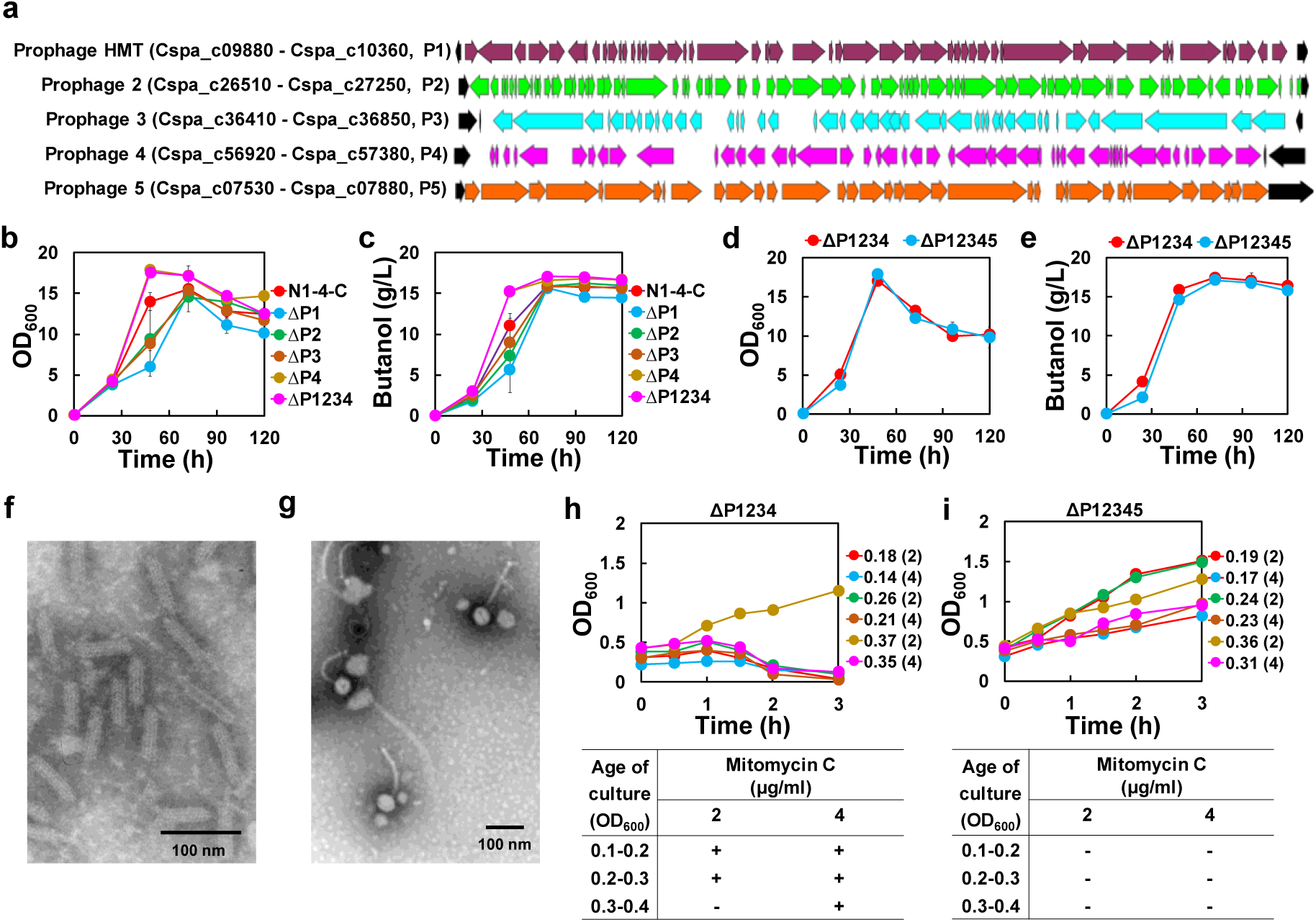
The deletion of the prophage genomes. (a) The gene cluster organization of the HM T prophage (P1) and another four putative prophages (P2-P5). (b) Comparison of the cell growth of prophage deleted mutants and the control N1-4-C strain. (c) Comparison of the butanol production in prophage deleted mutants and the control N1-4-C strain. (d) Comparison of the cell growth between ΔP1234 and ΔP12345. (e) Comparison of butanol production between ΔP1234 and ΔP12345; (f) Transmission election microscopy image of clostocin O; (g) Transmission election microscopy image of the HM T prophage particles; (h, i) Cell growth profiles ofΔP1234 and ΔP12345 with the induction (at various OD_600_) using mitomycin C at 2 or 4 µg/ml. “-” indicates that there was no cell lysis; “+” indicates that most of the cells were lysed. The value at the right side of the cell growth profile figure represents the actual OD_600_ value at which mitomycin C (with the applied concentration included in the parentheses) was added for the induction.

Thus, in a further step, we used ΔP1234 and ΔP12345 as the platform to be engineered for enhanced and more stable BA production. We deleted *nouG* and integrated *sadh*-*hydG* cluster in both ΔP1234 and ΔP12345, and obtained FJ-1200 and FJ-1300 correspondingly. The plasmid pMTL-P_adh_-*atf1-MinD* was transformed into FJ-1200 and FJ-1300, generating FJ-1201 and FJ-1301, respectively. Fermentation results showed that FJ-1201 and FJ-1301 produced 19.7 g/L and 19.4 g/L BA, respectively, which were both higher than FJ-308 (Fig. 6a & Table S3). BA production in both FJ-1201 and FJ-1301 could be completed within 48 h, resulting in a productivity of ∼0.41 g/L/h, which was significantly higher than 0.23 g/L/h in FJ-308. BA yield in FJ-1201 reached 0.26 g/g, which was also higher than FJ-308 (0.24 g/g). We further determined BA concentrations in the fermentation broth as 0.6 g/L and 0.5 g/L respectively for the fermentation with FJ-1201 and FJ-1301. Taken together, total BA production in FJ-1201 was 20.3 g/L, which was the highest level that has ever been reported in a microbial host. It is 2400-fold higher than the highest level that has been previously reported^22^, and also 14.8-fold higher than that by the very recently reported *C. diolis* strain^24^.

**Fig. 6.**
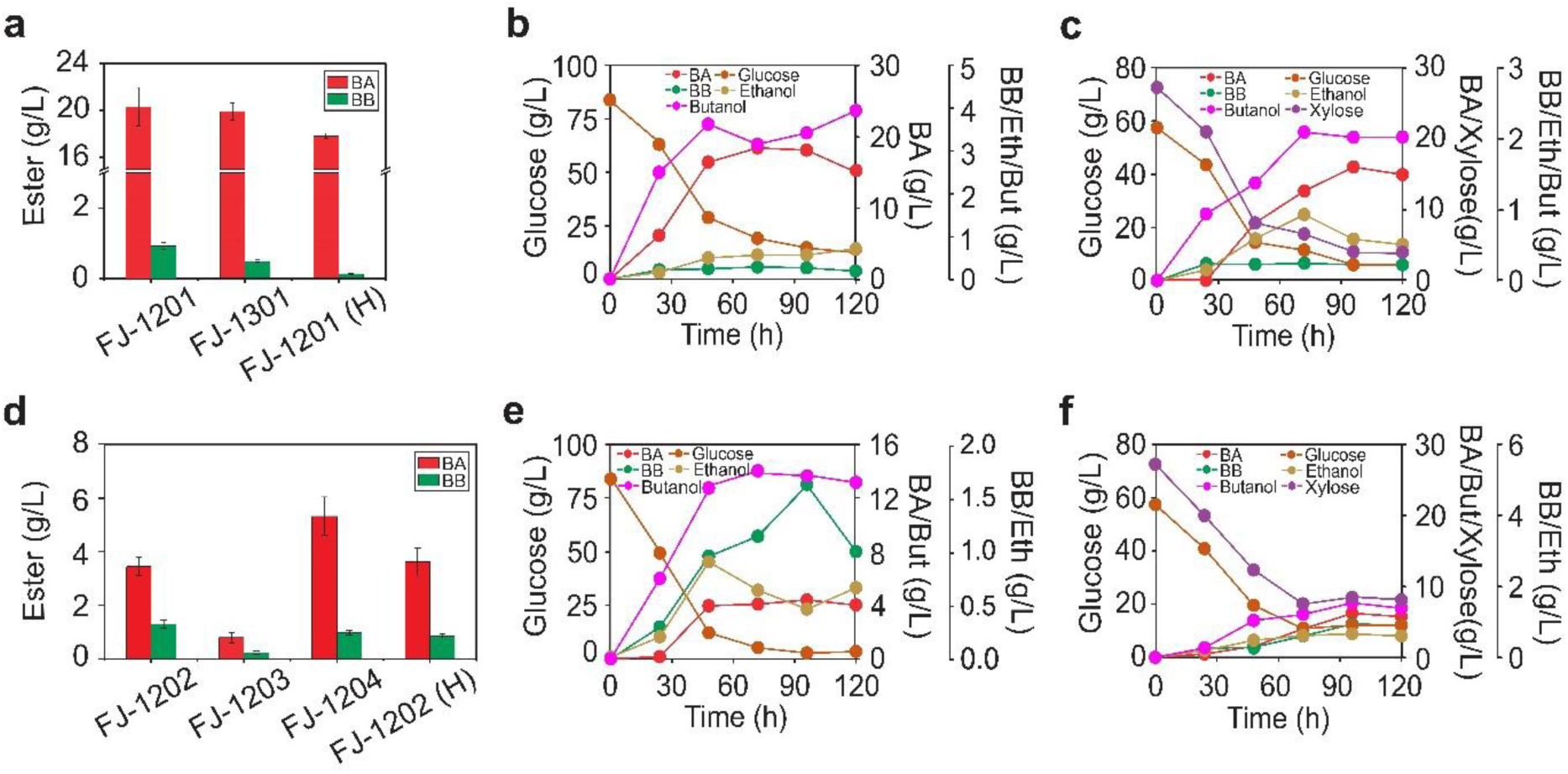
Fermentation results of the engineered strains for fatty acid ester production. (a) BA production in serum bottles using glucose (FJ-1201 and FJ-1301) or biomass hydrolysates (FJ-1201 (H)) as the substrate. (b) BA fermentation kinetics in FJ-1201 in 500-mL bioreactor using glucose as the substrate. (c) BA fermentation kinetics in FJ-1201 in 500-mL bioreactor using biomass hydrolysates as the substrate. (d) BB production in serum bottles using glucose (FJ-1202, FJ-1203, and FJ-1204) or biomass hydrolysates (FJ-1202 (H)) as the substrate. (e) BB fermentation kinetics in FJ-1202 in 500-mL bioreactor using glucose as the substrate. (f) BB fermentation kinetics in FJ-1202 in 500-mL bioreactor using biomass hydrolysates as the substrate. BA: butyl acetate; BB: butyl butyrate; Eth: ethanol; But: butanol.

Besides BA production, BB production in FJ-1201 also reached 0.9 g/L, which was significantly higher than in FJ-308 (0.01 g/L) and in *C. pasteurianum* J-5 (0.3 g/L) (Fig. 2b, Fig. 6a, & Table S3). This level (0.9 g/L) was 18.6-fold higher than the highest BB production that has been previously reported (0.05 g/L in *C. acetobutylicum*)^22^. All these results confirmed our hypothesis that the elimination of prophages would make more robust host strain for enhanced and more stable ester production.

### Expression of SAAT in FJ-1200 further enhanced BB production

As demonstrated in Fig. 2b, SAAT and EHT1 were more relevant for BB production. Therefore, to achieve higher BB production, we expressed *saat* and *eht1* in FJ-1200 and obtained FJ-1202 and FJ-1203, respectively. Fermentation showed that FJ-1202 and FJ-1203 produced 1.3 and 0.2 g/L BB, respectively (Fig.6d). Further, we added MinD C-tag to the SAAT and introduced the recombinant gene into FJ-1200 and obtained FJ-1204 for an attempt to further improve BB production as observed for BA production in FJ-308. However, BB production in FJ-1204 was only 1.0 g/L. Notwithstanding, 1.3 g/L BB obtained in FJ-1202 is 25.8-fold higher than the highest level that has been previously reported^22^.

### Ester production with biomass hydrolysates as substrate

Fermentations were carried out using biomass hydrolysates as the substrate. In the hydrolysates, besides sugars (57.4 g/L glucose and 27.2 g/L xylose) as carbon source, there were also nutrients converted from biomass (corn stover). Therefore, we tested the effect of organic nitrogen (yeast and tryptone) of various levels on ester production. Interestingly, results showed that the highest BA production of 17.5 g/L was achieved in FJ-1201 (in the extractant phase) without any exogenous nitrogen source supplemented (Table S4). In addition, 0.3 g/L BA was detected in aqueous phase, making a total BA production of 17.8 g/L in FJ-1201 (Fig. 6a & Table S4). Although this was slightly lower than when glucose was used as substrate (20.3 g/L), fermentation with hydrolysates did not need any supplementation of nutrients, which could significantly reduce production cost. We further performed fermentation in 500-mL bioreactor with pH controlled >5.0. BA production reached 16.0 g/L with hydrolysates and 18.0 g/L with glucose as substrate, both of which were lower than results from fermentation under the same conditions but with serum bottles (Figs. 6b & 6c). During bioreactor fermentation, we noticed very strong smell of BA around the reactor. We suspected that significant evaporation during fermentation with bioreactor resulted in the lower level of final BA titers as compared to fermentation with serum bottle, which was securely sealed with only minimum outlet for releasing gases. An improved bioprocess needs to be carefully designed for larger scale fermentation to minimize BA evaporation and enhance BA production and recovery.

Furthermore, we performed fermentation with FJ-1202 for BB production using hydrolysates in both serum bottle and 500-mL bioreactor. Results demonstrated that BB production in serum bottle from hydrolysates was 0.9 g/L (compared to 1.3 g/L when glucose used as substrate; Fig. 6d). BB production in bioreactor from hydrolysates reached 0.9 g/L compared to 1.6 g/L when glucose was used as substrate (Figs. 6e & 6f). The results were consistent with the case for BA production that lower-level BB was obtained when hydrolysates (compared to glucose) was used as substrate. However, interestingly, larger scale fermentation with bioreactor produced slightly higher level of BB than the fermentation under the same conditions with serum bottle, which was different from the case for BA production. This might be because BB is less evaporative (in bioreactor) than BA.

### Techno-economic analysis (TEA) for BA production from biomass hydrolysates

We performed a techno-economic analysis (TEA) to evaluate the economic competitiveness of BA production from corn stover at a process capacity of 2,500 MT wet corn stover (20% moisture) per day. The whole process was developed based on the previous process using the deacetylation and disk refining (DDR) pretreatment to produce corn stover hydrolysate^38^, which was the substrate used for our fermentation experiments to produce BA. The detailed process information is summarized in the supplementary materials. The process is composed of eight sections including feedstock handling, DDR pretreatment and hydrolysis, BA fermentation, product recovery (distillation), wastewater treatment, steam and electricity generation, utilities, and chemical and product storage (Fig. 7a). Fig. 7b shows the equipment cost distribution of each process sections, with a total installed equipment cost of $263 million. The fermentation, steam & electricity co-generation, and wastewater treatment contribute significant percentage to the total installed equipment cost, which aligns well with previous TEA models for chemical production from biomass via fermentation^39,40^. The total capital investment (TCI) is $472 million by taking consideration of additional direct cost, indirect cost as well as working capitals (Table S5). From the process model, 95.2 kg of BA can be produced from 1 MT of corn stover, meanwhile significant amounts of butanol (11.1 kg), isopropanol (15.5 kg) and surplus electricity (209 kWh) are produced as coproducts (Fig. 7c). The BA production cost was estimated to be $986/MT (Fig. 7d), which is much lower than the current BA market price ranging between $1,200 and $1,400 per MT in year 2019 (based on the quotes from the industry^41^), showing the highly economic competitiveness of BA production using our engineered strain. By looking into the cost breakdown, the corn stover feedstock cost contributes the most (38.2%) to the BA production cost, followed by other chemicals (22.3%) and capital deprecation (18.0%) and utilities (14.5%). Sensitivity analysis shows that corn stover price, BA yield, and BA titer are the most sensitive input parameters to the BA production cost (Fig. S10).

**Fig. 7.**
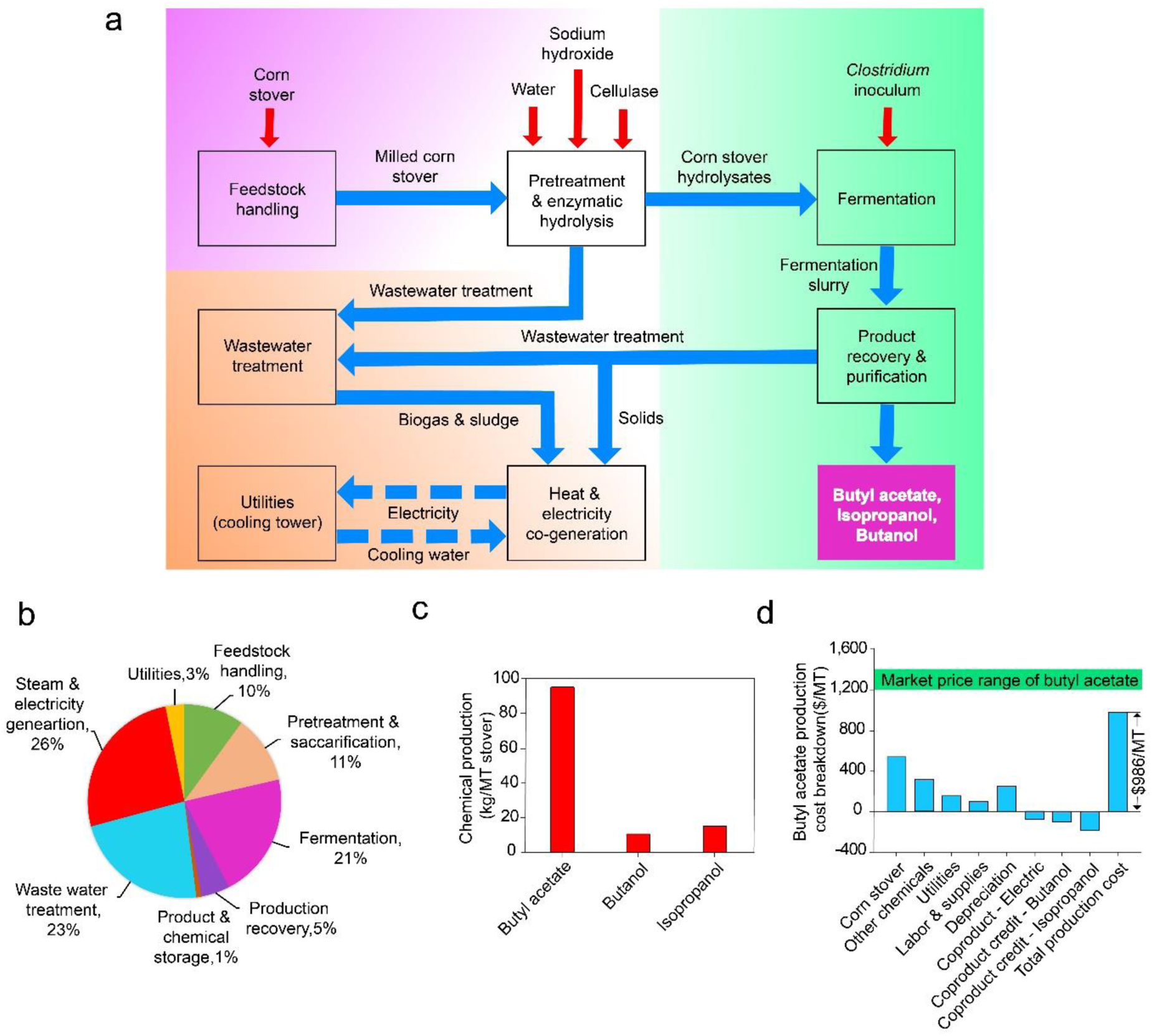
Techno-economic analysis of butyl acetate production from corn stover. a) process overview; b) total installed equipment cost; c) chemical production from each metric tonne (MT) of corn stover; d) butyl acetate production cost.

## Discussion

Although tremendous efforts have been invested on biofuel/biochemical research worldwide, very limited success has been achieved. A key bottleneck is that the microbial host is subject to endproduct toxicity and thus desirable production efficiency cannot be obtained^42^. Our central hypothesis was that metabolically engineering of microorganisms for high-value and easy-recoverable bioproduct production can help alleviate endproduct toxicity and thus high titer and productivity can be achieved, with which economically viable biofuel/biochemical production can be ultimately established. Here we tested this hypothesis by engineering solventogenic clostridia for high efficient ester (high-value and easy-recoverable) production. Our group have previously established versatile genome engineering tools for clostridia^11,14,27^, putting us at a strong position to perform this study.

Based on the systematic screening of host strains and enzymes as well as multiple rounds of rational metabolic engineering: enriching precursors (alcohols and acetyl-CoA) for ester production, dynamically expressing heterologous ester-production pathways, rationally organizing ester-synthesis enzymes, and improving strain robustness by eliminating putative prophages, we ultimately obtained strains for efficient production of esters in both synthetic fermentation medium and biomass hydrolysates. To the best of our knowledge, the production levels of BA and BB we achieved set up the new records. Overall, we demonstrated that clostridia are excellent platforms for valuable biofuel and biochemical production. The general principles that we demonstrated herein, including 1) selecting the most appropriate host for targeted bioproduction and 2) engineering the host for producing high value and easily recoverable products, are highly applicable to other relevant bioprocesses, and may result in breakthroughs in biofuel/biochemical production and general bioeconomy.

## Methods

### Microorganisms and cultivation conditions

All the strains and plasmids used in this study are listed in Table S1. *C. pasteurianum* ATCC 6013 and *C. saccharoperbutylacetonicum* N1-4 (HMT) (DSM 14923) were requested from American Type Culture Collection (ATCC) and Deutsche Sammlung von Mikroorganismen und Zellkulturen (DSMZ), respectively. *C. beijerinckii* NCIMB 8052 was provided by Dr. Hans P. Blaschek^14^. *C. tyrobutyricum* Δ*cat1*::*adhE1* and *C. tyrobutyricum* Δ*cat1*::*adhE2* are hyper-butanol producing mutants constructed in our lab^11^. All the clostridial strains were grown in an anaerobic chamber (N_2_-CO_2_-H_2_ with a volume ratio of 85:10:5) at 35 °C. Strains of *C. tyrobutyricum, C. saccharoperbutylacetonicum* and *C. beijerinckii* were cultivated using tryptone-glucose-yeast extract (TGY) medium^43^, while strains of *C. pasteurianum* were cultivated using 2×YTG medium^44^. When required, clarithromycin (Cla) or thiamphenicol (Tm) was supplemented into the medium at a final concentration of 30 µg/mL and 15 µg/mL, respectively. *E. coli* DH5α was used for routine plasmid propagation and maintenance. *E. coli* CA434 was used as the donor strain for plasmid conjugation for *C. tyrobutyricum*. Strains of *E. coli* were grown aerobically at 37 °C in Luria-Bertani (LB) medium supplemented with 100 µg/mL ampicillin (Amp), 50 μg/mL kanamycin (Kan) or 34 µg/mL chloramphenical (Cm) as needed.

### Plasmid construction

All the plasmids used in this study are listed in Table S1, and all the primers used in this study are listed in Table S2.

The plasmids pMTL82151 and pTJ1 were used as mother vectors for heterogeneous gene expression^45,46^. The promoter of the *cat1* gene (CTK_C06520) (P_*cat*_) and the promoter of the *thl* gene (CTK_C01450) (P_*thl*_) from *C. tyrobutyricum* ATCC 25755 were amplified and inserted into pMTL82151 at the *Eco*RI site, and the generated plasmids were named as pMTL82151-P_*cat*_ and pMTL82151-P_*thl*_, respectively. Promoters of the following gene, *pflA* (Cspa_c13710) (P_*pfl*_), *ald* (Cspa_c56880) (P_*ald*_), *adh* (Cspa_c04380) (P_*adh*_) and *bdh* (Cspa_c56790) (P_*bdh*_), all from *C. saccharoperbutylacetonicum* N1-4 (HMT) were amplified and inserted into pMTL82151 at the *Eco*RI site, and the generated plasmids were named as pMTL82151-P_*pfl*_, pMTL82151-P_*ald*_, pMTL82151-P_*adh*_ and pMTL82151-P_*bdh*_, respectively.

The *vaat* gene from *Fragaria vesca*, the *saat* gene from *F. ananassa* and the *atf1* gene from *S. cerevisiae* were amplified from plasmids pDL006, pDL001 and pDL004, respectively^3,21^.

The *atf1’* (the codon optimized *atf1* gene), *eht1* from *S. cerevisiae*^19^, *and lipaseB* from *Candida antarctica*^47^ *were all synthesized by GenScript (Piscataway, NJ, USA). The obtained gene fragments of vaat, saat, atf1, atf1’, eth1*, and *lipaseB* were inserted between the *Btg*ZI and *Eco*RI sites in pMTL82151-P_*cat*_, generating pMTL-P_*cat*_-*vaat*, pMTL-P_*cat*_-*saat*, pMTL-P_*cat*_-*atf1*, pMTL-P_cat_-*atf1*’, pMTL-P_cat_-*eht1*, and pMTL-P_cat_-*lipaseB*, respectively. The *atf1* gene was inserted between the *Btg*ZI and *Eco*RI sites in pMTL82151-P_*thl*_, generating pMTL-P_*thl*_-*atf1*.

The P_*cat*_ promoter and the gene fragments of *vaat, saat, atf1, eth1*, and *lipase* were amplified and ligated into the *Eco*RI site of pTJ1, generating pTJ1-P_*cat*_-*vaat*, pTJ1-P_*cat*_-*saat*, pTJ1-P_*cat*_-*atf1*, pTJ1-P_*cat*_-*eht1* and pTJ1-P_*cat*_-*lipaseB*, respectively. The *atf1* gene was inserted into the *Eco*RI site of pMTL82151-P_*pfl*_, pMTL82151-P_*ald*_, pMTL82151-P_*adh*_ and pMTL82151-P_*bdh*_, generating pMTL-P_*pfl*_-*atf1*, pMTL-P_*ald*_-*atf1*, pMTL-P_*adh*_-*atf1* and pMTL-P_*bdh*-_*aft1*, respectively.

DNA sequences of *CC-Di-A, CC-Di-B, MinD* and *pduA\** were synthesized by GenScript (Piscataway, NJ, USA). The MinD-tag was fused to the end of *atf1* with PCR and ligated into the *Eco*RI site of pMTL-P_*adh*_, generating pMTL-P_*adh*_-*atf1*-*MinD*. In addition, the MinD-tag was fused to the end of *saat* with PCR and inserted between the *Btg*ZI and *Eco*RI sites of pMTL-P_*cat*_, generating pMTL-P_*cat*_-*saat*-*MinD*.

The synthesized *CC-Di-A* fragment was ligated into the *Eco*RI site of pMTL-P_*adh*_, generating pMTL-P_*adh*_-CC-Di-A. The DNA fragments of *atf1* and *atf1*-*MinD* were amplified from pMTL-P_*adh*_-*atf1* and pMTL-P_*adh*_-*atf1*-*MinD* and then inserted into the *Eco*RI site of pMTL-P_*adh*_-CC-Di-A, obtaining pMTL-P_*adh*_-A-*atf1* and pMTL-P_*adh*_-A-*atf1*-*MinD*. The *CC-Di-B* sequence with the *nifJ* gene and *CC-Di-B* with the *bdhA* gene were subsequently inserted into the *Kpn*I site of pMTL-P_*adh*_-A-*atf1*, generating pMTL-P_*adh*_-A-*atf1*-B-*nifJ* and pMTL-P_*adh*_-A-*atf1-*B-*nifJ-*B-*bdhA*. The DNA fragments of *CC-Di-B*-*pduA*\*, *CC-Di-A*-*nifJ, CC-Di-A*-*bdhA* were inserted into the *Eco*RI site of pTJ1-P_*cat*_, generating pTJ1-P_*cat*_-B-*pduA*\*, pTJ1-P_*cat*_-B-*pduA*\*-A-*nifJ* and pTJ1-P_*cat*_-B-*pduA*\*-A-*nifJ*-A-*bdhA*, respectively.

For the gene deletion or integration in *C. saccharoperbutylacetonicum*, all the relevant plasmids were constructed based on pYW34, which carries the customized CRISPR-Cas9 system for genome editing in *C. saccharoperbutylacetonicum*^*14,27*^. The promoter *P*_*J23119*_ and the gRNA (with 20-nt guide sequence targeting on the specific gene) were amplified by two rounds of PCR with primers N-20nt/YW1342 and YW1339/YW1342 as described previously (N represents the targeted gene)^27^. The obtained fragment was then inserted into pYW34 (digested with *Btg*ZI and *Not*I) through Gibson Assembly, generating the intermediate vectors. For gene deletion, the fragment containing the two corresponding homology arms (∼500-bp for each) for deleting the specific gene through homologous recombination was amplified and inserted into the *Not*I site of the obtained intermediate vector as described above, generating pYW34-ΔN (N represents the targeted gene). For gene integration, the fragment containing the two corresponding homology arms (∼1000-bp for each), the promoter and the gene fragment to be integrated, was amplified and inserted into the *Not*I site of the obtained intermediate vector as described above, generating the final plasmid for gene integration purpose.

### Fermentation with glucose as the substrate

For the fermentation for ester production, the *C. pasteurianum* strain was cultivated in Biebl medium^48^ with 50 g/L glycerol as the carbon source at 35 °C in the anaerobic chamber. When the OD_600_ reached ∼0.8, the seed culture was inoculated at a ratio of 10% into 100 mL of the same medium in a 250-mL serum bottle and then cultivated at an agitation of 150 rpm and 30 °C (on a shaker incubator) for 72 h. The *C. beijerinckii* strain was cultivated in TGY medium until the OD_600_ reached ∼0.8. Then the seed culture was inoculated at a ratio of 5% into 100 mL P2 medium along with 60 g/L glucose and 1 g/L yeast extract in a 250-mL serum bottle. The fermentation was carried out at an agitation of 150 rpm and 37 °C for 72 h^43^. The *C. tyrobutyricum* strain was cultivated in RCM medium at 35 °C until the OD_600_ reached ∼1.5. Then the seed culture was inoculated at a ratio of 5% into 200 mL fermentation medium (containing: 50 g/L glucose, 5 g/L yeast extract, 5 g/L tryptone, 3 g/L (NH_4_)_2_SO_4_, 1.5 g/L K_2_HPO_4_, 0.6 g/L MgSO_4_·7H_2_O, 0.03 g/L FeSO_4_·7H_2_O, and 1 g/L L-cysteine) in a 500-mL bioreactor (GS-MFC, Shanghai Gu Xin biological technology Co., Shanghai, China) and the fermentation was carried out at an agitation of 150 rpm and 37 °C for 120 h with pH controlled >6.0^9^. The *C. saccharoperbutylacetonicum* strain was cultivated in TGY medium at 35 °C in the anaerobic chamber until the OD_600_ reached ∼0.8. Then the seed culture was inoculated at a ratio of 5% into 100 mL P2 medium along with 60 g/L glucose and 1 g/L yeast extract in a 250-mL serum bottle. The fermentation was carried out at an agitation of 150 rpm and 30 °C for 120 h^43^. For the fermentation at larger scales in bioreactors, it was carried out in a 500-mL fermenter (GS-MFC, Shanghai Gu Xin biological technology Co., Shanghai, China) with a working volume of 250 mL with pH controlled >5.0, at 50 rpm and 30 °C for 120 h. Samples were taken every 24 h for analysis.

For all fermentations in the serum bottle, a needle and hosepipe were connected to the top of bottle for releasing the gases produced during the fermentation. For all the fermentations for ester production, the extractant *n*-hexadecane was added into the fermentation with a ratio of 1:1 (volume of the extractant vs. volume of fermentation broth) for *in situ* ester extraction. The reported ester concentrations were the determined values in the extractant phase. All the fermentations were carried out in triplicate.

### Fermentation with biomass hydrolysates as the substrate

The biomass hydrolysates was kindly provided by Dr. Daniel Schell from National Renewable Energy Laboratory (NREL) which was generated from corn stover through the innovative ‘deacetylation and mechanical refining in a disc refiner (DDR)’ approach^49^. For the fermentation, the biomass hydrolysate was diluted and supplemented into the P2 medium as the carbon source (with final sugar concentrations of 57.4 g/L glucose and 27.2 g/L xylose). In addition, various concentrations of yeast extract (Y, g/L) and tryptone (T, g/L) were also added as the nitrogen source to evaluate their effects on the fermentation performance: 0Y+0T; 1Y+3T and 2Y+6T. The fermentation was carried out under the same conditions as described above at 100 mL working volume in a 250-mL serum bottle. All the fermentations were carried out in triplicate.

### Analytical methods

Concentrations of acetone, ethanol, butanol, acetic acid, butyric acid and glucose were measured using a high-performance liquid chromatography (HPLC, Agilent Technologies 1260 Infinity series, Santa Clara, CA) with a refractive index Detector (RID), equipped with an Aminex HPX-87H column (Bio-Rad Laboratories, Hercules, CA). The column was eluted with 5 mM H_2_SO_4_ at a flow rate of 0.6 mL/min at 25 °C. The concentration of the ester in the *n*-hexadecane phase was quantified using a gas chromatography-mass spectrometry (GC-MS, Agilent Technologies 6890N, Santa Clara, CA) equipped with an HP-5 column (60m×0.25 mm, 0.25 mm film thickness). Helium was used as the carrier gas. The initial temperature of the oven was set at 30 °C for 2 min, followed by a ramp of 10 °C/min to reach 300 °C, and a ramp of 2 °C/min to reach the final temperature of 320 °C, which was then held for 2 min. The detector was kept at 225 °C^9^.

## Authors’ contributions

J.F., Y.W. and I.B. designed the experiments. J.F., J.Z. (Jie Zhang), P.W., Y.G., Z.T.Z. and M.C. performed the experiments. Y.F. and H.H. performed the techno-economic analysis (TEA). J.F., Y.F., H.H. and Y.W. drafted the manuscript. P.J.B., Y.G., and J.Z. (Junping Zhou) contributed to improve the figures. M.C. and Z.S. contributed to the manuscript revision. All authors read and approved the manuscript.

## Data availability

The materials and data reported in this study are available upon reasonable request from the corresponding author.

## Acknowledgments

We thank Dr. Hans Blaschek (University of Illinois at Urbana-Champaign) for providing plasmids pYW34 and pTJ1, Dr. Nigel Minton (University of Nottingham) for providing pMTL series of plasmids, Dr. Cong T Trinh (University of Tennessee) for providing plasmids pDL001, pDL004 and pDL006, and Dr. Mike Young (Aberystwyth University, UK) for providing *E. coli* CA434. We thank Ms. Sheng Dong for constructing *C. pasteurianum* SD-1, and thank Drs. Michael Pyne and C. Perry Chou (University of Waterloo, Canada) for providing research materials and technical assistance for working on *C. pasteurianum*. We thank Dr. Daniel Schell (National Renewable Energy Laboratory) for providing the corn stover hydrolysates. We also thank Drs. Raymond P. Henry and Michael E. Miller (Auburn University) for allowing us to access their equipment for the phage preparation and TEM imaging.

## Author information

**Correspondence and requests for materials** should be addressed to H. H. or Y.W.

### Competing interests

Auburn University has filed a patent application covering the work described in this article*. The application names Y.W. and J.F. as inventors.

## Funding

This work was supported by the US Department of Energy’s Office of Energy Efficiency and Renewable Energy under Award DE-EE0008483 (Co-Optima), the Agriculture and Food Research Initiative Competitive Grant no. 2018-67021-27715 from the USDA National Institute of Food and Agriculture (NIFA), the Auburn University Intramural Grants Program (IGP), the USDA-NIFA Hatch project (ALA014-1017025), and the Alabama Agricultural Experiment Station.

## Supplementary information

*Supplementary Information*

Supplementary Information includes partial of the Methods, Results, Discussion and Supplementary Figs. 1–10 and Tables 1–8.

## Footnotes

* The initial provisional patent was filed on December 19, 2017.

## Notes

### Summary of Updates

The results for ethyl acetate production have been revised. Techno-economic analysis (TEA) has been added.

