## Supplementary materials for "Renewable Fatty Acid Ester Production in *Clostridium*"

[*]Corresponding author:

Haibo Huang,

### Methods

#### Plasmid transformation and mutant verification

Competent cells of *C. pasteurianum* were prepared following the protocol as reported by Pyne et al[1]. *C. pasteurianum* SD-1, an equivalent to *C. pasteurianum* Δ*cpaAIR* in which the *cpaAIR* gene (encoding the CpaAI Type II restriction endonuclease) was deleted and thus more efficient transformation was enabled[2], was used as the host strain. The overnight-grown seed culture was inoculated into 20 mL 2×YTG medium. When the $OD_{600}$ reached 0.3~0.4, sucrose and glycine were added to a final concentration of 0.4 M and 1.25%, respectively. When the $OD_{600}$ further reached 0.6~0.8, the cells were harvested by centrifugation at 4,200 g and 4 °C for 10 min. The cell pellets were resuspended in 5 mL of SMP buffer (270 mM sucrose, 1mM $MgCl_2$ and 7mM sodium phosphate, pH 6.5) and spinned down under the same conditions. The obtained cell pellets were then resuspended in 0.6 mL of SMP buffer. 1 μg of plasmid DNA was suspended with 20 μL of 2 mM Tris-HCl (pH 8.0) and then mixed with 550 μL competent cells and 30 μL 96% cold ethanol. The mixture was transferred to a pre-chilled 4 mm electroporation cuvette and incubated on ice for 5 min. Electroporation was then applied with a voltage of 1,800 V, capacitance of 25 μF and resistance of ∞ using a Gene Pulser Xcell electroporation system (Bio-Rad Laboratories, Hercules, CA). Afterwards, the culture was transferred into 2 mL of 2×YTG medium and recovered at 35 °C for 4 h. The culture was then spread onto the 2×YTGT agar plates (2×YTG agar plates containing 15 μg/mL of thiamphenicol) for the selection of the transformants.

Competent cells of *C. beijerinckii* were prepared following the procedure as described by Wang et al.[3]. Briefly, the overnight cell culture was inoculated into TGY medium with an inoculation ratio of 1%. When the $OD_{600}$ reached ~0.8, the cells were harvested by centrifugation at 4,200 g and 4 °C for 10 min. The cell pellets were washed with the same volume (as the original cell culture) of ice-cold 15% glycerol and centrifugated under the same condition for 10 min. The cell pellets were resuspended with 5% volume of ice-cold 15% glycerol. Then 400 μL competent cells and ~1.0 μg of plasmid DNA were mixed and transferred into a 2-mm pre-chilled electroporation cuvette and incubated on ice for 10 min. Electroporation was carried out at 2,000 V of voltage, 25 μF of capacitance and 200 Ω of resistance. Afterwards, the cells were transferred into 1.6 mL of TGY and incubated at 35 °C for 6-8 h for recovery. The culture was then spread onto TGYC agar plates (TGY agar plates containing 30 μg/mL of clarithromycin) for the selection of transformants.

The plasmid transformation through conjugation for *C. tyrobutyricum* was performed following the procedure as described by Zhang et al.[4]. The donor strain *E. coli* CA434 carrying the desired plasmid was cultivated in LB medium supplemented with Cm and Kan. 3 mL of overnight-cultured *E. coli* CA434 cells were centrifuged and washed for twice (with fresh LB medium) to remove the antibiotics. The obtained donor cells were then mixed with 0.4 mL of the overnight-cultured *C. tyrobutyricum* (grown in TGY medium). The cell mixture was spotted onto the TGY agar plate and incubated in the anaerobic chamber at 37 °C for conjugation. After 24 h of cultivation, the cell lawn on the plate was washed off using 1mL of TGY medium and then spread onto the TGY plate containing 15 μg/mL Tm and 250 μg/mL D-cycloserine (for eliminating the residual *E. coli* CA434 donor cells). The transformant colonies could be observed after 48-72 h of incubation in the anaerobic chamber.

Competent cells of *C. saccharoperbutylacetonicum* were prepared following the procedure as described by Herman et al., with slight modifications[5]. Briefly, the overnight-cultured cells were inoculated into fresh TGY medium. When the $OD_{600}$ of the cell culture reached ~0.8, cells were collected by centrifugation at 4,200 g and 22 °C (room temperature) for 10 min. Cell pellets were then washed with the same volume (as the original cell culture) of SMP buffer. The resuspension was centrifuged again under the same conditions as described above. The cell pellets were resuspended in 5% volume of SMP buffer. After that, the plasmid

(~1.0 μg) was mixed with 400 μL of competent cells and transferred into a 2 mm electroporation cuvette and incubated on ice for 30 min. Electroporation was then applied with a voltage of 1,000 V, capacitance of 25 μF and resistance of 300 Ω. Subsequently, the culture was transferred into 2 mL pre-warmed TGY medium and incubated at 35 °C for 2-3 h. After that, the culture was spread onto TGYC or TGYT (TGY agar plates containing 15 μg/mL of thiamphenicol) agar plates.

The positive mutants of *C. saccharoperbutylacetonicum* with desired gene deletion or integration were identified following our previously described procedures[6]. Briefly, the *C. saccharoperbutylacetonicum* transformants harboring the plasmid designed for the gene deletion or integration were incubated in TGYC liquid medium at 35 °C in the anaerobic chamber for about 24 h. The cell culture was then spread onto TGYLC plates (TGYC supplemented with 40 mM lactose). When colonies appeared on the plates, colony PCR (cPCR) was then carried out with the pair of primers of N-U/N-D (N represents the targeted gene name, U represents the upstream primer flanking the target locus, and D represents the downstream primer flanking the target locus) to verify the gene deletion or gene integration. The selected mutants were then subcultured in TGY medium for 3 to 5 generations to cure the plasmid for the gene deletion/integration[6]. The obtained plasmid-free and marker-free mutant strains were used for the following steps.

### Prophage induction and phage harvesting

The strain was grown in TGY medium for overnight. The cells were then transferred into fresh TGY prior to be expose to the inducing reagent mitomycin C or norfloxacin. Mitomycin C was added into the culture when the cell growth reached the desired $OD_{600}$ (0.1-0.2, 0.2-0.3, 0.3-0.4 or 0.4-0.5). The cells of some of the cultures grew too fast at the early stage for the induction purpose, and thus they were not induced under every above mentioned $OD_{600}$ conditions in this work. For the induction, generally mitomycin C at final concentrations of 2 μg/mL and 4 μg/mL was used[7]. Besides, 1 μg/mL and 3 μg/mL of mitomycin C were also tried for the induction in ΔP234. After 30 min of treatment at 35 °C, the cells were harvested via centrifugation at 4,000 g for 5 min and resuspended at the same volume of TGY fresh medium. Then the $OD_{600}$ was monitored carefully during the following 3-5 h.

The seed culture with the $OD_{600}$ of around 0.2, 0.4 and 0.6 was treated using norfloxacin at the final concentrations of 0.3, 1, 3, 6, 9 3 μg/mL at 35 °C for 24 h[8]. The final $OD_{600}$ was measured after the treatment.

The cells were collected via centrifugation at 4,000 g for 20 min, and then filtered through the 0.2 μm filter to obtain the supernatants. The supernatants were centrifugated at 20,000 rpm for 3h, and the obtained precipitation (containing the phages) was then resuspended by 1/30 volume of $ddH_2O$.

**Transmission election microscopy (TEM)**

20 μL prepared phage sample was applied to a mesh copper grid and settled for 2 min. Then the liquid was blotted off with a filter paper. The negative stain of 2% phosphotungstic acid (PTA) was applied to a mesh copper grid and settled for 30 s. Then the liquid was blotted off with a filter paper followed by air dry. The prepared samples were then used for TEM observation under a Zeiss EM10 transmission electron microscope (Carl Zeiss AG, Oberkochen, Germany) at an accelerating voltage of 60 kV.

**Techno-economic analysis (TEA)**

A comprehensive TEA model was developed to evaluate the economic feasibility of BA production from corn stover using the deacetylation and disk refining (DDR) pretreatment. The model originally developed to produce ethanol[9] was modified to produce BA by mainly substituting the fermentation and distillation unit operations. The processing capacity was set at 2,500 wet metric tonnes (MT, 20% moisture) of corn stover per day. The process was assumed to run 350 days (8,410 hours) per year; thus the annual corn stover consumption is 875,000 wet MT per year. The composition of corn stover was 35.05% cellulose, 19.53% hemicellulose, 15.76% lignin, 4.93% ash, 3.10% of protein, and 21.63% other solids on a dry basis[10]. All process was simulated using the software SuperPro Designer (Intelligen Inc., NJ).

The whole process can be divided into eight sections, including feedstock handling, DDR pretreatment and hydrolysis, fermentation, product recovery (distillation), wastewater treatment, steam and electricity cogeneration, utilities, and chemical and product storage. In the process, corn stover is milled at the pre-processing plant and delivered to the feed handling section from a uniform corn stover supply system[10]. Received corn stover is added with water to obtain a 25% solid slurry. The slurry is added with sodium hydroxide at a loading of 40 kg/MT dry corn stover, heated to 80 °C and held for 2 hours to remove acetyl groups from corn stover. The alkali-treated corn stover is then washed by using the same amount of added water, followed by dewatering using screw-type presses to remove excess water to attain 40% solids content for the subsequent disk refining and enzymatic hydrolysis. The parameters of disk refining are adapted from a previous optimization study using wet disk mills[9]. The electricity consumption of disk milling is 212 kWh per dry MT of corn stover. The details of the DDR process are described in Chen et al. 2015[9]. The pretreated corn stover is then cooled and sent to hydrolysis tanks where it is hydrolyzed into monomeric sugars by cellulase at 48 °C for 84 hr. The enzyme loading for the enzymatic hydrolysis is 19 mg protein/g cellulose according to a previous study[9]. After the enzymatic hydrolysis, 10% of the hydrolysate is split off to seed fermenters for production of seed culture, and the rest 90% of the hydrolysates is fermented into BA and coproducts in large fermenters at 32 °C, where hexadecane is added to the fermenter at 1:1 ratio (v/v) to extract BA from the fermentation broth. The fermentation takes 96 hours to convert all hydrolyzed sugars to BA and coproducts butanol and isopropanol. The key parameters for the conversion and fermentation yields are summarized in Table S6. The fermentation beer is then sent to the decanter to separate the extractant phase and the aqueous phase, which are separately pumped to the distillation section to recovery BA and coproducts butanol and isopropanol. The distillation stillage is press filtered to separate insoluble solids for combustion to produce steam and electricity, whereas the pressed filtrate is sent to the waste treatment section. The remaining sections of the model, including wastewater treatment, steam and electricity cogeneration, and utilities, inherited most of the original designs from the NREL process model[9, 10].

The energy and mass balance and flow rate information for the process were generated to determine the capital and operating costs. The purchased equipment costs were determined based on previous literature, particularly from Humbird et al. (2011)[10] and Chen et al. (2015)[9]. The cost of the product recovery (distillation) section was mainly determined by the embedded cost estimator of SuperPro Designer. The purchased equipment costs were scaled using the exponential scaling equation with exponents ranged between 0.5 and 0.8 depending on the type of equipment[10]. The equipment cost obtained in previous years is adjusted to the year of 2019 using the plant cost index from chemical engineering magazine. The total capital investment (TCI) was calculated as a sum of direct and indirect costs, which were determined based on the installed equipment costs. Direct costs included installed equipment cost, site development (9% of inside-battery-limits (ISBL) equipment cost), warehouse (4.0% of ISBL), and additional piping (4.5% of ISBL). Indirect cost is the sum of prorateable costs (10% of total direct cost (TDC)), field expenses (10% of TDC), home office and construction (20% of TDC), project contingency (10% of TDC), and other costs (10% of TDC). Working capital was assumed to be 5% of the fixed capital investment. A summary of the total operating cost is listed in Table S5. The total operating cost included both fixed and variable operating costs. Fixed operating costs include labor and various overhead items and variable operating costs include raw material costs, utility costs, and co-product credits. The detailed variable and fixed operating costs are summarized in Table S7.

The BA production cost was calculated based on the methods described in prior studies[11, 12]. For the process model, BA was assigned as the main product and butanol, isopropanol, and electricity were assigned as the coproducts, which were sold to generate coproduct credits. Sensitivity analyses were also performed at ± 20% variation range to evaluate the most influential variables on the BA production cost.

### Results

#### Evaluation of codon-optimization of *atf1* for BA synthesis

Different microorganisms have different codon usage preferences. The *atf1* gene was originally from *S. cerevisiae* and its genetic codon usage might not be preferable for the *C. saccharoperbutylacetonicum* host. Thus, a codon optimized *atf1* gene (designated as *atf'*) was synthesized and evaluated for potentially improved expression and BA production in the *C. saccharoperbutylacetonicum* host strain. However, fermentation results demonstrated that FJ-007 (carrying *atf1'* rather than *atf1*) actually generated slightly lower concentration of BA than FJ-004 (5.0 g/L vs. 5.5 g/L, Table S8). The result was unexpected, but not totally surprising. Similar cases have been reported previously where the original natural gene showed better efficiency than the codon-optimized counterpart for desirable biochemical production[13]. We speculate that the natural gene might be able to transcribe into more stable mRNA structure, and thus lead to higher translation level than the codon-optimized gene[13]. In addition, it has been recently reported that codon-optimized genes could bring about toxicity to the host cells[14].

#### Enhancement of acetyl-CoA availability to improve BA production

In the pathway, thiolase is the enzyme that coverts acetyl-CoA into acetoacetyl-CoA (Fig.

3a), and the attenuation of the thiolase activity might be able to repress the acetyl-CoA flux towards butanol (and other products) and lead to the accumulation of intracellular acetyl-CoA, which could be beneficial for enhanced BA production. Therefore, we attempted to delete the thiolase gene to further increase the acetyl-CoA availability. There are five annotated genes encoding thiolase in *C. saccharoperbutylacetonicum*: *Cspa_c06180*, *Cspa_c17310*, *Cspa_c20520*, *Cspa_c20890*, *Cspa_c50320*. After numerous attempts, we were only able to delete *Cspa_c20890* and *Cspa_c50320* individually in FJ-300, generating the mutant strains FJ-400 and FJ-500, respectively. By introducing the vector pMTL-cat-*atf1* into these two strains for BA production, FJ-401 and FJ-501 were obtained, respectively. Fermentation results indicated that BA production was actually slightly lower in FJ-401 (10.8 g/L) and FJ-501 (9.8 g/L) than that in FJ-301 (Fig. 3b). The result was unexpected. Acetoacetyl-CoA is one of the key metabolites for the cell; the repression for its biosynthesis might impair the cell metabolism and thus lead to decreased BA production.

**The elimination of prophages increased cell growth and butanol production**

During our fermentation process, we noticed that the performance for ester production of the strains was not very stable and could be varied from batch to batch. Our industrial collaborator also observed that *C. saccharoperbutylacetonicum* often had instable performance for ABE production in the continuous fermentation process (data not shown). It has been previously reported that the N1-4 (HMT) strain contains a temperate phage named HM T which could release from the chromosome even without induction[15]. In addition, the N1-4 (HMT) strain can produce a phage-like particle clostocin O with the induction of mitomycin C. We hypothesized that the instability of the fermentation with *C. saccharoperbutylacetonicum* might be related to the existence of prophages, and the deletion of these prophages and clostocin O encoding sequences would improve the stability of the strain and thus enable more stable and enhanced production of the desired endproduct (BA here). The online program PHAST was used to predict the prophage sequences in N1-4 (HMT)[16], with four possible prophage genomes were identified: the HM T prophage (renamed as P1) as well as three other putative prophages which were named as P2, P3 and P4 here (Fig. 5a). Besides, one additional incomplete prophage genome (without integrase gene) was found and named as P5.

We firstly constructed the mutant with single deletion of each prophage genome and generated the ΔP1, ΔP2, ΔP3 and ΔP4 strains. Because there are genes within the prophage genome possibly responsible for the normal cell metabolism, we also constructed the mutant with the deletion of only the integrase gene (without the integrase, the prophage cannot release from the chromosome), obtaining the ΔNP1, ΔNP2, ΔNP3 and ΔNP4 strains. In addition, we also constructed the mutant ΔP1234 (with the deletion of all four prophage genomes) and ΔNP1234 (with the deletion of all four integrase genes).

Fermentations were first conducted in the serum bottle to investigate the effects of the elimination of prophages on butanol production in the mutant strains. As shown in Figs. 5b, 5c & S3, ΔP4 and ΔP1234 showed increased cell growth and butanol production, while all the other eight mutants did not demonstrate significance difference in terms of the cell growth and solvent production compared to the mother strain. ΔP4 and ΔP1234 reached the maximum $OD_{600}$ of 17.9 and 17.6, which were 15.5% and 13.5% higher than that of the control N1-4-C strain, respectively. The butanol production in ΔP4 and ΔP1234 reached 17.1 and 16.8 g/L respectively, which were also higher than that of the control N1-4-C strain (16.0 g/L).

To further study the individual prophage, we constructed the triple-deletion mutants ΔP234, ΔP134, ΔP124, ΔP123. Phage induction experiments of ΔP234, ΔP134, ΔP124, ΔP123 and ΔP1234 with mitomycin C revealed that all the mutants exhibited cell lysis (Fig. S4). Transmission election microscopy (TEM) results indicated that all the mutants produced the tail-like particles, which showed likely the same appearance as clostocin O as reported previously[7] (Fig. S5). We were not able to observe the HM T phage in the supernatant of ΔP234. It might be because there were too many clostocin O particles in the view which made it difficult to observe the HM T phage.

Based on the above results, we tentatively concluded that P5 might be responsible for the production of clostocin O. To verify this hypothesis and obtain a more robust strain for bioproduction, P5 was deleted in ΔP234, ΔP134, ΔP124, ΔP123 and ΔP1234, obtaining ΔP2345, ΔP1345, ΔP1245, ΔP1235 and ΔP12345. The induction experiments indicated that the cell lysis was detected in ΔP2345 with the addition of 4 μg/mL of mitomycin C at the $OD_{600}$ of 0.2-0.5 (Fig. S6), while no cell lysis was detected in any other mutants at any conditions with the treatment using mitomycin C or norfloxacin (data not shown). Furthermore, after induction, phage-like particles were only observed in the supernatant of ΔP2345 (Fig. 5g & Fig. S7) and they were likely the HM T phages. However, the phage image was different from what was described before[17]. It was more like HM 7 (a head with a long tail), rather than HM 1 (a head with multiple short tails). It is worthwhile to mention that this is the first time that an image of the HM T phage has been reported.

After the deletion of P5, no clostocin O particle was observed in the supernatant of ΔP12345, which confirmed that P5 indeed encoded clostocin O. In addition, as mentioned above, no cell lysis was observed in ΔP12345 with induction (Fig. 5i & Fig. S6), suggesting that ΔP12345 could be a more stable platform to be engineered for enhanced ester production. On the other hand, we showed above that ΔP1234 grew faster and produced more butanol than the control N1-4-C strain. Therefore, we further compared the fermentation performance of ΔP1234 vs ΔP12345 in both serum bottles and bioreactors (Figs. 5d, 5e, S8, & S9). Results showed that the further deletion of P5 in ΔP12345 did not result in significant difference in cell growth or butanol production when compared to ΔP1234; actually the butanol production in ΔP12345 was slightly lower than in ΔP1234.

### Discussion

We firstly set out to screen the host strains and ester synthesis genes for specific ester production. With the combination of five clostridial strains (*C. saccharoperbutylacetonicum* N1-4-C, *C. pasteurianum* ester SD-1, *C. beijerinckii* 8052, *C. tyrobutyricum cat1::adhE1* and *cat1::adhE2*) and five ester synthesis genes (*vaat*, *saat*, *atf1*, *eht1* and *lipaseB*), we obtained very promising results. Most of the engineered strains could produce EA, BA and BB at the same time (Fig. 2), and some of the strains could also produce small amount of EB. *C. saccharoperbutylacetonicum* FJ-004 produced 5.5 g/L BA, which was the highest BA production level that has ever been reported[18]. *C. pasteurianum* J-5 produced 0.3 g/L BB, which was also significantly higher than the previously reported level of 0.05 g/L in an engineered *C. acetobutylicum* strain[18, 19]. The results confirmed our hypothesis that solventogenic clostridia are outstanding platforms to be engineered for ester production. Because the BA production in FJ-004 was significantly higher than the production levels of other esters, we decided to focus on further improving BA production in *C. saccharoperbutylacetonicum* through systematic metabolic engineering.

Butanol and acetyl-CoA are the two precursors for BA synthesis. The enhancement of the intracellular pool of these two precursors in the host could help improve BA production. We thus firstly deleted *nuoG* to save NADH and improve butanol and thus BA production. Our fermentation results showed that FJ-101 with the deletion of *nuoG* had increased BA production to 7.8 g/L. However, there was still 7.6 g/L butanol remaining at the end of fermentation with FJ-101; it was thus reasonable to speculate that the availability of acetyl-CoA was the bottleneck for further improving BA production. We tried two strategies to improve intracellular acetyl-CoA availability. One was for the enhanced 'regeneration' of acetyl-CoA, and the other was for 'blocking' the pathway that consumes acetyl-CoA. Comparatively, the former seemed a better strategy. By introducing a heterologous isopropanol synthesis pathway to promote the 'regeneration' of intracellular acetyl-CoA, the FJ-301 strain could produce up to 12.9 g/L BA (Fig. 3b).

The dynamic expression of the heterologous pathway to be synchronous with the production of the precursors could be highly beneficial for the production of the target bioproduct. On the other hand, the imbalance of intracellular metabolism and the accumulation of toxic precursors would harm the cells and lead to decreased production of the target product. Previously, synthetic regulatory tools have been developed to dynamically control the gene expression and result in enhanced production of the targeted bioproducts. For example, Zhang et al., (2012) developed a dynamic sensor-regulator system (DSRS) by engineering a hybrid fatty acid/acyl-CoA-regulated promoter[20]. The promoter can sense the level of the precursors for the synthesis of fatty acid ethyl ester and thus regulate the gene expression level in response to the physiological state of the cell, therefore increased the target product level. However, it needs tremendous works to screen the best responsive engineered promoter. It is well known that the intracellular metabolism especially the solvent production in clostridia is strictly regulated by the cells[21]. We hypothesized that the appropriate regulation of the BA synthesis enzyme using the native promoter of the host strain could achieve the similar effect as DSRS. In this work, four native promoters associated with BA precursors formation were selected and evaluated to control the *atf1* gene expression (Fig. 3c). Results indicated that the *atf1* gene controlled by the $P_{adh}$ promoter showed 10.5% increase in BA production compared with the control FJ-301 strain (Fig. 3d). The $P_{adh}$ promoter is responsible for the ethanol and butanol synthesis. The synchronous expression of alcohol dehydrogenase and ATF1 remarkably increased BA production.

Spatial organization of the enzymes associated with BA synthesis is another strategy that we employed to enhance BA production. The cross-link of the enzymes associated with BA synthesis or anchoring these enzymes onto a synthetic scaffold (PduA*) was not able to improve the BA production; while anchoring the ATF1 enzyme to the cell membrane by adding a MinD C-tag to the C-terminus of the enzyme led to significantly increased BA production. The obtained FJ-308 produced 16.4 g/L BA, which was 20% more than that in FJ-304 (Fig. 4). The attachment of ATF1 to the cell membrane facilitated the excretion of BA from the cells, which could mitigate the intracellular toxicity caused by BA and meanwhile boost the BA synthesis.

During the fermentation, the performance of the strain for BA production was not stable, and remarkable cell lysis was also observed at the end of the fermentation. We speculated that the instability of the strain could be because of the prophages existing in the chromosome of *C. saccharoperbutylacetonicum*. Based on analysis, we identified four putative prophages P1-P4 and one incomplete prophage genome P5 in the genome of *C. saccharoperbutylacetonicum* N1-4 (HMT). P5 was demonstrated to be responsible for the synthesis of clostocin O (Figs. 5f & S5). Ultimately, we obtained two mutant strains ΔP1234 and ΔP12345, both of which grew faster and produced more butanol than the wild type strain (Figs. 5b-e, S3, S8, & S9). Thus, we further constructed the BA-producing strains FJ-1201 and FJ-1301 respectively based on ΔP1234 and ΔP12345. Fermentation results demonstrated that FJ-1201 could produce 20.3 g/L BA (Fig. 6a & Table S3), which was the highest production level of BA that has ever been reported in any microbial biocatalyst host[18]. The BA yield in FJ-1201 reached 0.26 g/g, which was also significantly higher than the initial BA-producing strain FJ-004 (0.07 g/g). Thus, the deletion of prophages from *C. saccharoperbutylacetonicum* could not only increase the cell growth (and stability) but also the production of the desired endproducts (butanol or BA).

In addition, we also noticed that the BB production in FJ-1201 reached 0.9 g/L, which was significantly higher than that in *C. pasteurianum* J-5 (0.3 g/L, the highest BB production level based on our initial screening of the strains and enzymes for ester production) (Fig. 6a & Table S3). We further overexpressed *saat* (instead of *atf1*) in the strain, and obtained the FJ-

1202 strain in which the BB production reached unprecedented 1.3 g/L (Fig. 6d).

Both the BA-producing *C. saccharoperbutylacetonicum* FJ-1201 and the BB-producing *C. saccharoperbutylacetonicum* FJ-1202 performed well when biomass hydrolysates was used as the substrate for the fermentation. FJ-1201 could produce 17.8 g/L BA and FJ-1202 could produce 0.9 g/L BB from biomass hydrolysates (with no need to supplement any exogenous nitrogen source). Although these levels were slightly lower than when glucose was used as the substrate for the fermentation with the same strain, the operation eliminated the requirement of yeast extract and tryptone and thus would significantly decrease the cost of the bioprocess for fatty acid ester production.

### Table S1 Strains and plasmids used in this study

| Strains and plasmids | Relevant genotype and characteristics | Source |
|---|---|---|
| **Strains** | | |
| *C. saccharoperbutylacetonicum* N1-4 (HMT) | DSM 14923 (= ATCC 27021), wild-type strain | DSMZ |
| *C. tyrobutyricum* Δ*cat1::adhE1* | ATCC 25755, *cat1::adhE1* | [4] |
| *C. tyrobutyricum* Δ*cat1::adhE2* | ATCC 25755, *cat1::adhE2* | [4] |
| *C. pasteurianum* SD-1 | ATCC 6013 with the deletion of *cpaAI* gene | Lab stock |
| *Clostridium beijerinckii* NCIMB 8052 | Wild-type strain | [22] |
| *C. saccharoperbutylacetonicum* N1-4-C | ΔCsp_135p (with the endogenous plasmid Csp_135p eliminated) | [23] |
| *C. saccharoperbutylacetonicum* FJ-001 | N1-4-C harboring pMTL82151 | This study |
| *C. saccharoperbutylacetonicum* FJ-002 | N1-4-C harboring pMTL-$P_{cat}$-*vaat* | This study |
| *C. saccharoperbutylacetonicum* FJ-003 | N1-4-C harboring pMTL-$P_{cat}$-*saat* | This study |
| *C. saccharoperbutylacetonicum* FJ-004 | N1-4-C harboring pMTL-$P_{cat}$-*atf1* | This study |
| *C. saccharoperbutylacetonicum* FJ-005 | N1-4-C harboring pMTL-$P_{cat}$-*eht1* | This study |
| *C. saccharoperbutylacetonicum* FJ-006 | N1-4-C harboring pMTL-$P_{cat}$-*lipaseB* | This study |
| *C. saccharoperbutylacetonicum* FJ-007 | N1-4-C harboring pMTL-$P_{cat}$-*atf1'* | This study |
| *C. saccharoperbutylacetonicum* FJ-008 | N1-4-C harboring pMTL-$P_{thl}$-*atf1* | This study |
| *C. saccharoperbutylacetonicum* FJ-100 | N1-4-C, Δ*nuoG* | This study |
| *C. saccharoperbutylacetonicum* FJ-101 | FJ-100 harboring pMTL-$P_{cat}$-*atf1* | This study |
| *C. saccharoperbutylacetonicum* FJ-200 | FJ-100 with $P_{thl}$-*sadh* integrated at *Cspa_c21540* site | This study |
| *C. saccharoperbutylacetonicum* FJ-201 | FJ-200 harboring pMTL-$P_{cat}$-*atf1* | This study |
| *C. saccharoperbutylacetonicum* FJ-300 | FJ-100 with $P_{thl}$-*sadh-hydG* integrated at *Cspa_c21540* site | This study |
| *C. saccharoperbutylacetonicum* FJ-301 | FJ-300 harboring pMTL-$P_{cat}$-*atf1* | This study |
| *C. saccharoperbutylacetonicum* FJ-302 | FJ-300 harboring pMTL-$P_{pfl}$-*atf1* | This study |
| *C. saccharoperbutylacetonicum* | FJ-300 harboring pMTL-$P_{ald}$-*atf1* | This study |

| | | |
|---|---|---|
| FJ-303 | | |
| *C. saccharoperbutylacetonicum* FJ-304 | FJ-300 harboring pMTL-$P_{adh}$-*atf1* | This study |
| *C. saccharoperbutylacetonicum* FJ-305 | FJ-300 harboring pMTL-$P_{bdh}$-*atf1* | This study |
| *C. saccharoperbutylacetonicum* FJ-306 | FJ-300 harboring pMTL-$P_{adh}$-A-*atf1* | This study |
| *C. saccharoperbutylacetonicum* FJ-307 | FJ-300 harboring pMTL-$P_{adh}$-A-*atf1*-MinD | This study |
| *C. saccharoperbutylacetonicum* FJ-308 | FJ-300 harboring pMTL-$P_{adh}$-*atf1*-MinD | This study |
| *C. saccharoperbutylacetonicum* FJ-309 | FJ-300 harboring pMTL-$P_{adh}$-A-*atf1*-B-*nifJ* | This study |
| *C. saccharoperbutylacetonicum* FJ-310 | FJ-300 harboring pMTL-$P_{adh}$-A-*atf1*-B-*nifJ*-B-*bdhA* | This study |
| *C. saccharoperbutylacetonicum* FJ-311 | FJ-300 harboring pTJ1-$P_{cat}$-B-*pduA** and pMTL-$P_{adh}$-A-*atf1* | This study |
| *C. saccharoperbutylacetonicum* FJ-312 | FJ-300 harboring pTJ1-$P_{cat}$-B-*pduA**-A-*nifJ and* pMTL-$P_{adh}$-A-*atf1* | This study |
| *C. saccharoperbutylacetonicum* FJ-313 | FJ-300 harboring pTJ1-$P_{cat}$-B-*pduA**-A-*nifJ*-A-*bdhA* and pMTL-$P_{adh}$-A-*atf1* | This study |
| *C. saccharoperbutylacetonicum* FJ-400 | FJ-300 derivative, Δ*Cspa_c20890* | This study |
| *C. saccharoperbutylacetonicum* FJ-401 | FJ-400 harboring pMTL-$P_{cat}$-*atf1* | This study |
| *C. saccharoperbutylacetonicum* FJ-500 | FJ-300 derivative, Δ*Cspa_c50320* | This study |
| *C. saccharoperbutylacetonicum* FJ-501 | FJ-500 harboring pMTL-$P_{cat}$-*atf1* | This study |
| *C. saccharoperbutylacetonicum* FJ-1100 | ΔP1234 derivative with the deletion of *nuoG* | This study |
| *C. saccharoperbutylacetonicum* FJ-1200 | ΔP1234 derivative with the insertion of $P_{thl}$-*sadh-hydG* | This study |
| *C. saccharoperbutylacetonicum* FJ-1201 | FJ-1200 harboring pMTL-$P_{adh}$-*atf1*-MinD | This study |
| *C. saccharoperbutylacetonicum* FJ-1300 | ΔP12345 derivative with the deletion of *nuoG* and insertion of $P_{thl}$-*sadh-hydG* | This study |
| *C. saccharoperbutylacetonicum* FJ-1301 | FJ-1300 harboring pMTL-$P_{adh}$-*atf1*-MinD | This study |
| *C. saccharoperbutylacetonicum* FJ-1202 | FJ-1200 harboring pMTL-$P_{cat}$-*saat* | This study |
| *C. saccharoperbutylacetonicum* FJ-1203 | FJ-1200 harboring pMTL-$P_{cat}$-*eht1* | This study |

| | | |
|---|---|---|
| *C. saccharoperbutylacetonicum* FJ-1204 | FJ-1200 harboring pMTL-P$_{cat}$-*saat*-MinD | This study |
| *C. saccharoperbutylacetonicum* ΔP1 | N1-4-C strain with the deletion of prophage HM T (P1) | This study |
| *C. saccharoperbutylacetonicum* ΔP2 | N1-4-C strain with the deletion of putative phage P2 | This study |
| *C. saccharoperbutylacetonicum* ΔP3 | N1-4-C strain with the deletion of putative phage P3 | This study |
| *C. saccharoperbutylacetonicum* ΔP4 | N1-4-C strain with the deletion of putative phage P4 | This study |
| *C. saccharoperbutylacetonicum* ΔNP1 | N1-4-C strain with the deletion of *Cspa_c09880* (NP1) gene | This study |
| *C. saccharoperbutylacetonicum* ΔNP2 | N1-4-C strain with the deletion of *Cspa_c26510* (NP2) gene | This study |
| *C. saccharoperbutylacetonicum* ΔNP3 | N1-4-C strain with the deletion of *Cspa_c36800* (NP3) gene | This study |
| *C. saccharoperbutylacetonicum* ΔNP4 | N1-4-C strain with the deletion of *Cspa_c57380* (NP4) gene | This study |
| *C. saccharoperbutylacetonicum* ΔNP1234 | N1-4-C strain with the deletion of *Cspa_c09880*, *Cspa_c26510*, *Cspa_c36800* and *Cspa_c57380* | This study |
| *C. saccharoperbutylacetonicum* ΔP123 | N1-4-C strain with the deletion of putative prophages P1, P2 and P3 | This study |
| *C. saccharoperbutylacetonicum* ΔP124 | N1-4-C strain with the deletion of putative prophages P1, P2 and P4 | This study |
| *C. saccharoperbutylacetonicum* ΔP134 | N1-4-C strain with the deletion of putative prophages P1, P3 and P4 | This study |
| *C. saccharoperbutylacetonicum* ΔP234 | N1-4-C strain with the deletion of putative prophages P2, P3 and P4 | This study |
| *C. saccharoperbutylacetonicum* ΔP1234 | N1-4-C strain with the deletion of putative prophages P1, P2, P3 and P4 | This study |
| *C. saccharoperbutylacetonicum* ΔP2345 | N1-4-C strain with the deletion of putative prophages P2, P3, P4 and P5 | This study |
| *C. saccharoperbutylacetonicum* ΔP1345 | N1-4-C strain with the deletion of putative prophages P1, P3, P4 and P5 | This study |
| *C. saccharoperbutylacetonicum* ΔP1245 | N1-4-C strain with the deletion of putative prophages P1, P2, P4 and P5 | This study |
| *C. saccharoperbutylacetonicum* ΔP1235 | N1-4-C strain with the deletion of putative prophages P1, P2, P3 and P5 | This study |
| *C. saccharoperbutylacetonicum* ΔP12345 | N1-4-C strain with the deletion of putative prophages P1, P2, P3, P4 and P5 | This study |
| *C. pasteurianum* J-1 | SD-1 harboring pMTL82151 | This study |
| *C. pasteurianum* J-2 | SD-1 harboring pMTL-P$_{cat}$-*vaat* | This study |
| *C. pasteurianum* J-3 | SD-1 harboring pMTL-P$_{cat}$-*saat* | This study |

| | | |
|---|---|---|
| *C. pasteurianum* J-4 | SD-1 harboring pMTL-P$_{cat}$-*atf1* | This study |
| *C. pasteurianum* J-5 | SD-1 harboring pMTL-P$_{cat}$-*eht1* | This study |
| *C. pasteurianum* J-6 | SD-1 harboring pMTL-P$_{cat}$-*lipaseB* | This study |
| *Clostridium beijerinckii* F-1 | NCIMB 8052 harboring pTJ1 | This study |
| *Clostridium beijerinckii* F-2 | NCIMB 8052 harboring pTJ1-P$_{cat}$-*vaat* | This study |
| *Clostridium beijerinckii* F-3 | NCIMB 8052 harboring pTJ1-P$_{cat}$-*saat* | This study |
| *Clostridium beijerinckii* F-4 | NCIMB 8052 harboring pTJ1-P$_{cat}$-*atf1* | This study |
| *Clostridium beijerinckii* F-5 | NCIMB 8052 harboring pTJ1-P$_{cat}$-*eht1* | This study |
| *Clostridium beijerinckii* F-6 | NCIMB 8052 harboring pTJ1-P$_{cat}$-*lipaseB* | This study |
| *C. tyrobutyricum* JZ-1 | Δ*cat1::adhE1* harboring pMTL82152 | This study |
| *C. tyrobutyricum* JZ-2 | Δ*cat1::adhE1* harboring pMTL-P$_{cat}$-*vaat* | This study |
| *C. tyrobutyricum* JZ-3 | Δ*cat1::adhE1* harboring pMTL-P$_{cat}$-*saat* | This study |
| *C. tyrobutyricum* JZ-4 | Δ*cat1::adhE1* harboring pMTL-P$_{cat}$-*atf1* | This study |
| *C. tyrobutyricum* JZ-5 | Δ*cat1::adhE1* harboring pMTL-P$_{cat}$-*eht1* | This study |
| *C. tyrobutyricum* JZ-6 | Δ*cat1::adhE1* harboring pMTL-P$_{cat}$-*lipaseB* | This study |
| *C. tyrobutyricum* JZ-7 | Δ*cat1::adhE2* harboring pMTL82152 | This study |
| *C. tyrobutyricum* JZ-8 | Δ*cat1::adhE2* harboring pMTL-P$_{cat}$-*vaat* | This study |
| *C. tyrobutyricum* JZ-9 | Δ*cat1::adhE2* harboring pMTL-P$_{cat}$-*saat* | This study |
| *C. tyrobutyricum* JZ-10 | Δ*cat1::adhE2* harboring pMTL-P$_{cat}$-*atf1* | This stud |
| *C. tyrobutyricum* JZ-11 | Δ*cat1::adhE2* harboring pMTL-P$_{cat}$-*eht1* | This study |
| *C. tyrobutyricum* JZ-12 | Δ*cat1::adhE2* harboring pMTL-P$_{cat}$-*lipaseB* | This study |
| *E. coli* CA434 | hsd20(rB-, mB-), recA13, rpsL20, leu, proA2, with IncPb conjugative plasmid R702 | [24] |
| *E. coli* DH5α | F$^-$, φ80d*lacZ*ΔM1, Δ(*lacZYA-argF*)U169, *deoR*, *recA*1, *endA*1, *hsdR*17(r$_k^-$, m$_k^+$), *phoA*, *supE*44, λ$^-$ *thi*-1, *gyrA*96, *relA*1 | NEB |
| **Plasmids** | | |
| pYW34 | CAK *ori*, Amp$^r$, Erm$^r$, Plac::Cas9, gRNA | [3] |
| pMTL82151 | pBP1 *ori*, *catP*, ColE1, *tra* | [25] |
| pYW34-Δ*nuoG* | Derivative of pYW34 for *nuoG* deletion | This study |
| pYW34-*sadh* | Derivative of pYW34 for P$_{thl}$-*sadh* integration | [26] |
| pYW34-*sadh-hydG* | Derivative of pYW34 for P$_{thl}$-*sadh-hydG* integration | [26] |
| pYW34-Δ*Cspa_c06180* | Derivative of pYW34 for *Cspa_c06180* deletion | This study |
| pYW34-Δ*Cspa_c17310* | Derivative of pYW34 for *Cspa_c17310* deletion | This study |
| pYW34-Δ*Cspa_c20520* | Derivative of pYW34 for *Cspa_c20520* deletion | This study |
| pYW34-Δ*Cspa_c20890* | Derivative of pYW34 for *Cspa_c20890* deletion | This study |
| pYW34-Δ*Cspa_c50320* | Derivative of pYW34 for *Cspa_c50320* deletion | This study |
| pYW34-ΔP1 | Derivative of pYW34 for P1 phage deletion | This study |
| pYW34-ΔP2 | Derivative of pYW34 for P2 phage deletion | This study |
| pYW34-ΔP3 | Derivative of pYW34 for P3 phage deletion | This study |
| pYW34-ΔP4 | Derivative of pYW34 for P4 phage deletion | This study |
| pYW34-ΔP5 | Derivative of pYW34 for P5 phage deletion | This study |
| pYW34-ΔNP1 | Derivative of pYW34 for *Cspa_c09880* deletion | This study |
| pYW34-ΔNP2 | Derivative of pYW34 for *Cspa_c26510* deletion | This study |

| | | |
|---|---|---|
| pYW34-ΔNP3 | Derivative of pYW34 for *Cspa_c36800* deletion | This study |
| pYW34-ΔNP4 | Derivative of pYW34 for *Cspa_c57380* deletion | This study |
| pMTL82151-$P_{cat}$ | =pJZ98-P*cat1*, pMTL82151 derivative, containing $P_{cat}$ for gene overexpression purpose | [4] |
| pMTL82151-$P_{thl}$ | pMTL82151 derivative, containing $P_{thl}$ for gene overexpression purpose | Lab stock |
| pMTL82151-$P_{pfl}$ | pMTL82151 derivative, containing $P_{pfl}$ for gene overexpression purpose | This study |
| pMTL82151-$P_{bld}$ | pMTL82151 derivative, containing $P_{bld}$ for gene overexpression purpose | This study |
| pMTL82151-$P_{adh}$ | pMTL82151 derivative, containing $P_{adh}$ for gene overexpression purpose | This study |
| pMTL82151-$P_{bdh}$ | pMTL82151 derivative, containing $P_{bdh}$ for gene overexpression purpose | This study |
| pMTL-$P_{cat}$-*vaat* | pMTL82151-$P_{cat}$ derivative, with *vaat* inserted downstream of $P_{cat}$ | This study |
| pMTL-$P_{cat}$-*saat* | pMTL82151-$P_{cat}$ derivative, with *saat* inserted downstream of $P_{cat}$ | This study |
| pMTL-$P_{cat}$-*atf1* | pMTL82151-$P_{cat}$ derivative, with *atf1* inserted downstream of $P_{cat}$ | This study |
| pMTL-$P_{cat}$-*eht1* | pMTL82151-$P_{cat}$ derivative, with *eht1* inserted downstream of $P_{cat}$ | This study |
| pMTL-$P_{cat}$-*lipaseB* | pMTL82151-$P_{cat}$ derivative, with *lipaseB* inserted downstream of $P_{cat}$ | This study |
| pMTL-$P_{cat}$-*atf1'* | pMTL82151-$P_{cat}$ derivative, with *atf1'* inserted downstream of Pcat | This study |
| pMTL-$P_{thl}$-*atf1* | pMTL82151-$P_{thl}$ derivative, with *atf1* inserted downstream of $P_{thl}$ | This study |
| pMTL-$P_{pfl}$-*atf1* | pMTL82151-$P_{pfl}$ derivative, with *atf1* inserted downstream of $P_{pfl}$ | This study |
| pMTL-$P_{bld}$-*atf1* | pMTL82151-$P_{bld}$ derivative, with *atf1* inserted downstream of $P_{bld}$ | This study |
| pMTL-$P_{adh}$-*atf1* | pMTL82151-$P_{adh}$ derivative, with *atf1* inserted downstream of $P_{adh}$ | This study |
| pMTL-$P_{bdh}$-*atf1* | pMTL82151-$P_{bdh}$ derivative, with *atf1* inserted downstream of $P_{bdh}$ | This study |
| pMTL-$P_{adh}$-CC-Di-A | pMTL82151-$P_{adh}$ derivative, CC-Di-A inserted downstream of $P_{adh}$ | This study |
| pMTL-$P_{adh}$-*atf1*-MinD | pMTL82151-$P_{adh}$ derivative, with *atf1*-MinD inserted downstream of $P_{adh}$ | This study |
| pMTL-$P_{cat}$-*saat*-MinD | pMTL82151-$P_{cat}$ derivative, with *saat*-MinD inserted downstream of $P_{cat}$ | This study |
| pMTL-$P_{adh}$-A-*atf1* | pMTL82151-$P_{adh}$ derivative, with CC-Di-A-*atf1* | This study |

| | | |
|---|---|---|
| | fragment inserted downstream of $P_{adh}$ | |
| pMTL-$P_{adh}$-A-*atf1*-MinD | pMTL82151-$P_{adh}$ derivative, with CC-Di-A-*atf1*-MinD fragment inserted downstream of $P_{adh}$ | This study |
| pMTL-$P_{adh}$-A-*atf1*-B-*nifJ* | pMTL-$P_{adh}$-A-*atf1* derivative, with CC-Di-B-*nifJ* fragment inserted downstream of the *atf1* gene | This study |
| pMTL-$P_{adh}$-A-*atf1*-B-*nifJ*-B-*bdhA* | pMTL-$P_{adh}$-A-*atf1*-B-*nifJ* derivative, with CC-Di-B-*bdhA* fragment inserted downstream of the *nifJ* gene | This study |
| pMTL-$P_{adh}$-A-*atf1*-MinD-B-*nifJ* | pMTL-$P_{adh}$-A-*atf1*-MinD derivative, with CC-Di-B-*nifJ* fragment inserted downstream of the *atf1*-MinD gene | This study |
| pMTL-$P_{adh}$-A-*atf1*-MinD-B-*nifJ*-B-*bdhA* | pMTL-$P_{adh}$-A-*atf1*-MinD-B-*nifJ* derivative, with CC-Di-B-*bdhA* fragment inserted downstream of the *atf1*-MinD gene | This study |
| pTJ1-$P_{cat}$-B-*pduA** | pTJ1-$P_{cat}$ derivative with the insertion of CC-Di-B-*pduA** | This study |
| pTJ1-$P_{cat}$-B-*pduA**-A-*nifJ* | pTJ1-$P_{cat}$-B-*pduA** derivative with the insertion of CC-Di-A-*nifJ* | This study |
| pTJ1-$P_{cat}$-B-*pduA**-A-*nifJ*-A-*bdhA* | pTJ1-$P_{cat}$-B-*pduA**-A-*nifJ* derivative with the insertion of CC-Di-A-*bdhA* | This study |
| pTJ1 | pYL102E derivative, $Amp^r$, $Erm^r$ | [22] |
| pTJ1-$P_{cat}$ | pTJ1 derivative, with the insertion of $P_{cat}$ promoter | This study |
| pTJ1-$P_{cat}$-*vaat* | pTJ1 derivative, with the insertion of $P_{cat}$-*vaat* | This study |
| pTJ1-$P_{cat}$-*saat* | pTJ1 derivative, with the insertion of $P_{cat}$-*saat* | This study |
| pTJ1-$P_{cat}$-*atf1* | pTJ1 derivative, with the insertion of $P_{cat}$-*atf1* | This study |
| pTJ1-$P_{cat}$-*eht1* | pTJ1 derivative, with the insertion of $P_{cat}$-*eht1* | This study |
| pTJ1-$P_{cat}$-*lipaseB* | pTJ1 derivative, with the insertion of $P_{cat}$-*lipaseB* | This study |
| pDL004 | pETite* carrying *atf1* from *S. cerevisiae* | [27] |
| pDL001 | pETite* carrying *saat* from *Fragaria ananassa* | [28] |
| pDL006 | pETite* carrying *vaat* from *F. vesca* | [27] |

**Table S2 Primers used in this study**

| **Primers** | **Sequence (5' to 3')** | **Purpose** |
|---|---|---|
| nuoG-UF | TGATATGACTAATAATTAGCGGCCGCTAGTATTTTCAGGAATTTCTCCAAT | pYW34-Δ*nuoG* |
| nuoG-UR | TTGAAAGACATTAAGAGGGGCAACGATTAACTATGATAAATG | |
| nuoG-DF | TTAATCGTTGCCCCTCTTAATGTCTTTCAAACTAAATACC | |
| nuoG-DR | ACTAGTAACCATCACACTGTAAGATAACCTTTGAATTGTTACGC | |
| nuoG-U | ATTGCTTGAATTTCGTCTTCAAAAC | |
| nuoG-D | TAGATAACATGATATCCAAGGTCGC | |
| nuoG-20nt | GCTCAGTCCTAGGTATAATGCTAGCTTTACTTCATCTTTTGAAGCGTTTTAGAGCTAGAAATAGCAAG | |
| YW1139 | GTCATAAACTTGCTCAACTGATATGATCTTCCATAACTTAAC | gRNA amplification |
| YW1142 | ATATAGGTAATCGCTTTCATAAGGGCCCGATCGGTCTTGCCTTGCTCGTCG | |
| Cspa_c06180-UF | TGATATGACTAATAATTAGCGGCCGCCCCCATTAGATACTAAAGAAACTCA | pYW34-Δ*Cspa_c06180* |
| Cspa_c06180-UR | CTAAACTCTTTCAACAACTACGTCTCTCATGTTTGACCTC | |
| Cspa_c06180-DF | ATGAGAGACGTAGTTGTTGAAAGAGTTTAGTATACAAGTT | |
| Cspa_c06180-DR | ACTAGTAACCATCACACTGGGTTTGTAAAAGCAGTTTATTCATC | |
| Cspa_c06180-U | GTTTTTGAGAGATTAGGTGGAAAGG | |
| Cspa_c06180-D | TCCCTACTGCATAGATTCCTGCATT | |
| Cspa_c06180-20nt | GCTCAGTCCTAGGTATAATGCTAGCTCAAAAGTAAATGTTAATGGGTTTTAGAGCTAGAAATAGCAAG | |
| Cspa_c17310-UF | TGATATGACTAATAATTAGCGGCCGCCAGACAGTGGCCTTATCAATAGGAG | pYW34-Δ*Cspa_c17310* |
| Cspa_c17310-UR | TTATAATCTTTCTACAACTACTTCTTTCATTTATGCTCCT | |
| Cspa_c17310-DF | ATGAAAGAAGTAGTTGTAGAAAGATTATAATAATAGATAA | |
| Cspa_c17310-DR | ACTAGTAACCATCACACTGCCTCTTTTTTCTTTTTCATTGCTTC | |
| Cspa_c17310-U | TCAGGGACAGCAGTTCATGTAGTAA | |
| Cspa_c17310-D | TTATGGTAAAGCCTAATACAAACCC | |
| Cspa_c17310-20nt | GCTCAGTCCTAGGTATAATGCTAGCAAGAAAGTTAATGTAAGTGGGTTTTAGAGCTAGAAATAGCAAG | |
| Cspa_c20520-UF | TGATATGACTAATAATTAGCGGCCGCAAGACATTAAAGATTCTGCAACAGC | pYW34-Δ*Cspa_c20520* |
| Cspa_c20520-UR | TTATTCTCTTTCAACTACTACATCTTTCATTTTTTATTCCTC | |
| Cspa_c20520-DF | ATGAAAGATGTAGTAGTTGAAAGAGAATAATTGAATTTAA | |
| Cspa_c20520-DR | ACTAGTAACCATCACACTGATGCAAGATACACAAGCTATAGGAT | |
| Cspa_c20520-U | TTTTATGTCATAGCAATTGAGGGAA | |

| | | |
|---|---|---|
| Cspa_c20520-D | TTGTTCTTTATATTTTTCATTGGGC | |
| Cspa_c20520-20nt | GCTCAGTCCTAGGTATAATGCTAGCCTTGCAACCTTGTGTATTGGGTTTTAGAGCTAGAAATAGCAAG | |
| Cspa_c20890-UF | TGATATGACTAATAATTAGCGGCCGCAATAACTGATGTAGCCAGTGCAACT | pYW34-Δ*Cspa_c20890* |
| Cspa_c20890-UR | TCAATTGCATCTTTCTACTACTTCTTTCATTTTTAACTCC | |
| Cspa_c20890-DF | ATGAAAGAAGTAGTAGAAAGATGCAATTGATAGAGGAATG | |
| Cspa_c20890-DR | ACTAGTAACCATCACACTGAATTTTCTTACCGCTACAAAGATCA | |
| Cspa_c20890-U | AATATTTGCTGAACTTGATGACGTA | |
| Cspa_c20890-D | ACCATTTAATACATAATGGTCACCT | |
| Cspa_c20890-20nt | GCTCAGTCCTAGGTATAATGCTAGCTGTTAAATGATGCAAGATGGGTTTTAGAGCTAGAAATAGCAAG | |
| Cspa_c50320-UF | TGATATGACTAATAATTAGCGGCCGCTGACAATAATGTGTTTAAATTTCCT | pYW34-Δ*Cspa_c50320* |
| Cspa_c50320-UR | GTGGAAAGTGTATATATTAAAAGGAGCTAAGGATGAACTAT | |
| Cspa_c50320-DF | TTAGCTCCTTTTAATATATACACTTTCCACTTATAAAACT | |
| Cspa_c50320-DR | ACTAGTAACCATCACACTGCTATTTTAGGATATGTTGATAATGC | |
| Cspa_c50320-U | TTTCATTGATTTTATCCATCTTCTT | |
| Cspa_c50320-D | TTAATCCTATTTATTCAGATGAAGA | |
| Cspa_c50320-20nt | GCTCAGTCCTAGGTATAATGCTAGCAACTGTGCTACTTGAAAATAGTTTTAGAGCTAGAAATAGCAAG | |
| P1-UF | TGATATGACTAATAATTAGCATCAACCAGGGAATTTATAAGAAGC | pYW34-ΔP1 |
| P1-UR | AGCTGCTAATTTAGTATCAGTCTTCATTTAATGGAATTTACATTAA | |
| P1-DF | TAAATGAAGACTGATACTAAATTAGCAGCTGAAAATGAAAGT | |
| P1-DR | ACTAGTAACCATCACACTGGCCCAAAATTCCTTTAACTTGCTCGTA | |
| P1-U | TATTAGTGCAAAATGGCTTTATGAC | |
| P1-D | GCATCTGAATCCATTATTTTAGTTT | |
| P1-20nt | GCTCAGTCCTAGGTATAATGCTAGCGATCATATATTGCTAAACATGTTTTAGAGCTAGAAATAGCAAG | |
| P2-UF | TGATATGACTAATAATTAGCGAAGAGAAAGAAAGCAAATATTACC | pYW34-ΔP2 |
| P2-UR | ATAGCTCCTATCTTTGTGTTAAAAATGTGTTAATAATAAA | |
| P2-DF | TTAACACATTTTTAACACAAAGATAGGAGCTATATATGAAA | |
| P2-DR | ACTAGTAACCATCACACTGGCTTTAACTTCGATCCTCTTAGCACCT | |
| P2-U | CCAAGTGAAAAAATTGAATTTGATA | |

| | | |
|---|---|---|
| P2-D | AATTTTTATTAAGACCCTGCCGCTA | |
| P2-20nt | GCTCAGTCCTAGGTATAATGCTAGCGGAGCTGCAACAACTA TAAAGTTTTAGAGCTAGAAATAGCAAG | |
| P3-UF | TGATATGACTAATAATTAGC GCTTAATATTGTCTATTGATACCGA | pYW34-ΔP3 |
| P3-UR | GAATAAAAGATGATTAGATATGCTCGAATTTGATTATTATA A | |
| P3-DF | AATTCGAGCATATCTAATCATCTTTTATTCTATACACATTA | |
| P3-DR | ACTAGTAACCATCACACTGGCTTATAAATCTTTATGGGGC AAGAGG | |
| P3-U | TTTTCCAAATAATTTGCTTAATCCT | |
| P3-D | CAAGACAACTAATTATCGAAAAACA | |
| P3-20nt | GCTCAGTCCTAGGTATAATGCTAGCTCACCTCCATATTTTCT TCTGTTTTAGAGCTAGAAATAGCAAG | |
| P4-UF | TGATATGACTAATAATTAGCCAGAGCTTTGTGTAATTGATG TAAC | pYW34-ΔP4 |
| P4-UR | TATATTATCAAGCAAATAACCCGTAAGTAACAATTAAGCT | |
| P4-DF | TTACTTACGGGTTATTTGCTTGATAATATATTTAAATATA | |
| P4-DR | ACTAGTAACCATCACACTGGCCTAATGAAGCCAATATGGA GAAACT | |
| P4-U | ATGCAACATACCGGAAAAGGTAAAC | |
| P4-D | AATAGATAGGTTGCAAAAGGAAATG | |
| P4-20nt | GCTCAGTCCTAGGTATAATGCTAGCTCTCTTAAATCACTCAT TATGTTTTAGAGCTAGAAATAGCAAG | |
| P5-UF | TGATATGACTAATAATTAGCATTTGTATGAAGGTAGCGGTT GAGG | pYW34-ΔP5 |
| P5-UR | TAAGATAGTTTAGTTTCCTTTACAAATTAATGTATTTTTTAA | |
| P5-DF | TTAATTTGTAAAGGAAACTAAACTATCTTAGAAGACAGAC | |
| P5-DR | ACTAGTAACCATCACACTGGCCTTAGGAATGATATTCGAA AAAGCA | |
| P5-U | TCTTTCCTTATAAAGAGTATCGTTTCTAA | |
| P5-D | CAATTAAAGTATGAACAAAAGCATGG | |
| P5-20nt | GCTCAGTCCTAGGTATAATGCTAGCTTCAAAATTACAAGCTT CATGTTTTAGAGCTAGAAATAGCAAG | |
| Cspa_c09880-UF | TGATATGACTAATAATTAGCAATTTCAAGCATAATGGTTAT TTCA | pYW34-ΔNP1 |
| Cspa_c09880-UR | ATGAAAGCAGCTATTAAAGGCGCTCTTTAATGTAGCGCCC | |
| Cspa_c09880-DF | TTAAAGAGCGCCTTTAATAGCTGCTTTCATAGTATACCTCC | |
| Cspa_c09880-DR | ACTAGTAACCATCACACTGGCTCTTAATCAGTCTTCATTTAA TGGA | |
| Cspa_c09880-U | GGAGATGCTTTTTATGATAAAGAAA | |
| Cspa_c09880-D | ATATCATTAATACTGATAATGGCGG | |
| Cspa_c09880-20nt | GCTCAGTCCTAGGTATAATGCTAGCCATTAACCAGTCTTGG | |

| | | |
|---|---|---|
| | TGCAGTTTTAGAGCTAGAAATAGCAAG | |
| Cspa_c26510-UF | TGATATGACTAATAATTAGCGAAAGCCTCAGGATGGAAAGATAGT | pYW34-ΔNP2 |
| Cspa_c26510-UR | ATGGCAAATAAAACTTTAAAAATGTGTTAATAATAAAAATAGC | |
| Cspa_c26510-DF | TTAACACATTTTTAAAGTTTTATTTGCCATAAAATCATCC | |
| Cspa_c26510-DR | ACTAGTAACCATCACACTGGCAGAAAACAGAAAGGATTAACTCAAC | |
| Cspa_c26510-U | GCTGTGCTGGTATCTAAAATAAAGG | |
| Cspa_c26510-D | AGGTGAAATTATTGTGTTAGGTGA | |
| Cspa_c26510-20nt | GCTCAGTCCTAGGTATAATGCTAGCGGAATAGTTACTTTTAAGCAGTTTTAGAGCTAGAAATAGCAAG | |
| Cspa_c36800-UF | TGATATGACTAATAATTAGCAGCTGACAAATTAAAAGTTAGTGAA | pYW34-ΔNP3 |
| Cspa_c36800-UR | CTAATTTAGCAACGCAGTCTTTACAGCCATAAGTAAAACC | |
| Cspa_c36800-DF | ATGGCTGTAAAGACTGCGTTGCTAAATTAGAAATATAAAC | |
| Cspa_c36800-DR | ACTAGTAACCATCACACTGGCGGAATGTTTATACCAGTCCTATTCA | |
| Cspa_c36800-U | AGAAAAATACAGAAATGATCTTCAA | |
| Cspa_c36800-D | GGTACTTAATTTCTCTTTGGTTTC | |
| Cspa_c36800-20nt | GCTCAGTCCTAGGTATAATGCTAGCTGATATAGATTATGATAATAGTTTTAGAGCTAGAAATAGCAAG | |
| Cspa_c57380-UF | TGATATGACTAATAATTAGCAATTTCTGAACATTTTAAAGGAAGA | pYW34-ΔNP4 |
| Cspa_c57380-UR | TTTAAATATATTATCCTTATAACTTGCCATAAATACCACC | |
| Cspa_c57380-DF | ATGGCAAGTTATAAGGATAATATATTTAAATATAAACCTC | |
| Cspa_c57380-DR | ACTAGTAACCATCACACTGGCCTAAGGAAGAAAACAAAATAGCCAA | |
| Cspa_c57380-U | TACATAAATTCTTTCCAGAAGGGAT | |
| Cspa_c57380-D | GCCCAATTAAAAAGTGGAGTAGAAG | |
| Cspa_c57380-20nt | GCTCAGTCCTAGGTATAATGCTAGCATATTTAAATGTTACCCACCGTTTTAGAGCTAGAAATAGCAAG | |
| pMTL-KZR | AGATCTCCATGGACGCGTGACGTCG | |
| $P_{bld}$-F | TCCATATGACCATGATTACGTTAAATAAATTTCCTTAAAAAACAATAT | pMTL82151-$P_{bld}$ |
| $P_{bld}$-R | ATCCCCGGGTACCGAGCTCGAATTCGACTAATCCTCCTTATGATTTAAAA | |
| $P_{bdh}$-F | TCCATATGACCATGATTACGGTTAAGCCACTACCAGGATCAATCA | pMTL82151-$P_{bdh}$ |
| $P_{bdh}$-R | ATCCCCGGGTACCGAGCTCGAATTCTTCTTTTCCTCCTCTTACACACGCA | |
| $P_{adh}$-F | TCCATATGACCATGATTACGACAGGAGATAGCAAATACAGAGCTA | pMTL82151-$P_{adh}$ |

| | | |
|---|---|---|
| $P_{adh}$-R | ATCCCCGGGTACCGAGCTCGAATTCAATTTTAATAACCCTCCTCAAAATA | |
| $P_{pfl}$-F | TCCATATGACCATGATTACGTTAGATCCTAAAAAATATGGCGTTA | pMTL82151-$P_{pfl}$ |
| $P_{pfl}$-R | ATCCCCGGGTACCGAGCTCGAATTCAAATTAACCACCTCCCAAATTGAAA | |
| vaat-F | ATATAATTTATGAAAGGGTGGTTTTTATGGAGAAAATTGAGGTCAGT | pMTL-$P_{cat}$-*vaat* |
| vaat-R | CGACTCTAGAGGATCCCCGGGTACCGAATTCTCAATATCTTGAAATTAGCGTC | |
| saat-F | ATATAATTTATGAAAGGGTGGTTTTTATGGAGAAAATTGAGGTCAGT | pMTL-$P_{cat}$-*saat* |
| saat-R | CGACTCTAGAGGATCCCCGGGTACCGAATTCTTAAATTAAGGTCTTTGGAGATGC | |
| atf1-F | ATATAATTTATGAAAGGGTGGTTTTTATGAATGAAATCGATGAGAAAAATCA | pMTL-$P_{cat}$-*atf1* |
| atf1-R | CGACTCTAGAGGATCCCCGGGTACCGAATTCCTAAGGGCCTAAAAGGAGAGCTTTGT | |
| eht1-F | ATATAATTTATGAAAGGGTGGTTTTTATGTCAGAAGTATCTAAGTGG | pMTL-$P_{cat}$-*eht1* |
| eht1-R | CGACTCTAGAGGATCCCCGGGTACCGAATTCTTATACAACCAACTCGTCAAAC | |
| lipaseB-F | ATATAATTTATGAAAGGGTGGTTTTTATGAGAAAAGTATTTTTAAGATC | pMTL-$P_{cat}$-*lipaseB* |
| lipaseB-R | CGACTCTAGAGGATCCCCGGGTACCGAATTCCTAAGGAGTAACAATTCCTGAG | |
| atf1’-F | ATATAATTTATGAAAGGGTGGTTTTTATGAACGAGATCGACGAGAAAAACC | pMTL-$P_{cat}$-*atf1* |
| atf1’-R | CGACTCTAGAGGATCCCCGGGTACCGAATTCTTACGGGCCTAACAGCAGCGC | |
| atf1-thl-F | ATATAATTTATGAAAGGGTGGTTTTTATGAATGAAATCGATGAGAAAAATCA | pMTL-$P_{thl}$-*atf1* |
| atf1-thl-R | CGACTCTAGAGGATCCCCGGGTACCGAATTCCTAAGGGCCTAAAAGGAGAGCTTTGT | |
| atf1-bld-F | AATCATAAGGAGGATTAGTCATGAATGAAATCGATGAGAAAAATCA | pMTL82151-$P_{bld}$-*atf1* |
| atf1-bdh-F | GTGTAAGAGGAGGAAAAGAAATGAATGAAATCGATGAGAAAAATCA | pMTL82151-$P_{bdh}$-*atf1* |
| atf1-adh-F | TGAGGAGGGTTATTAAAATTATGAATGAAATCGATGAGAAAAATCA | pMTL82151-$P_{adh}$-*atf1* |
| atf1-pfl-F | ATTTGGGAGGTGGTTAATTTATGAATGAAATCGATGAGAAAAATCA | pMTL82151-$P_{pfl}$-*atf1* |
| atf1-promoter-R | ATCCCCGGGTACCGAGCTCGAATTCCTAAGGGCCTAAAAGG | |

| | | |
|---|---|---|
| | AGAGCTTTGT | |
| atf1-MinD-R1 | AGCCATCATCCCTTTGTTTTGTTCCTCCAAGACTTGCAACGGCGATCCCGAACCACTAGTAGGGCCTAAAAGGAGAGCTT | Adding MinD-tag to *atf1* 3’ end |
| atf1-MinD-R2 | GATCCCCGGGTACCGAGCTCTTAGGAACGTACACCGAAAAATGATTTGATTTT | |
| saat-MinD-R1 | GCCATCATCCCTTTGTTTTGTTCCTCCAAGACTTGCAACGGCGATCCCGAACCACTAGTAATTAAGGTCTTTGGAGATGC | Adding MinD-tag to *saat* 3’ end |
| saat-MinD-R2 | CTAGAGGATCCCCGGGTACCGAGCTCTTAGGAACGTACACCGAAAAATGATTTGATTTTAGCCATCATCCCTTTGTTTTG | |
| adh-A-F | TGAGGAGGGTTATTAAAATTATGGGTAGCGAAATTGCCGCGCTGG | CC-Di-A amplification |
| adh-A-R | ATTTTTCTCATCGATTTCATTCATATGGCTGCCGCGCGGCACCA | |
| B-nifJ-F1 | ACTGAAAAAGAAAAACGCTGCGCTGAAACAGAAAATTGCCGCGCTGAAACAGATGAGAAAAATGAAAACTATGGATG | CC-Di-B-*nifJ* amplification |
| B-nifJ-F2 | TTTAGGCCCTTAGGAGCTCGAAAGAGGGGTCAACGCGATGGGTAGCAAAATCGCCGCACTGAAAAAGAAAAACGCTGCG | |
| B-nifJ-R | TCGACTCTAGAGGATCCCCGGGTACCCTATTGTTGATTAGCTAACTTCTTG | |
| B-bdhA-F1 | ACTGAAAAAGAAAAACGCTGCGCTGAAACAGAAAATTGCCGCGCTGAAACAGATGATGAGATTTACATTACCAAGAG | CC-Di-B-*bdhA* amplification |
| B-bdhA-F2 | GTTAGCTAATCAACAATAGGAGGAGGGTTATTAAAATTATGGGTAGCAAAATCGCCGCACTGAAAAAGAAAAACGCTGCG | |
| B-bdhA-R | TCGACTCTAGAGGATCCCCGGGTACCTTAAAAATCAACCTTATTTCCATAA | |
| pMTL-TJ1-F | TTGTAAAACGACGGCCAGTGGTAGACTTTAAGGATGGAACCTTTG | Amplification of fragments from pMTL82151-based plasmids |
| pMTL-TJ1-R | TTCTAACGCGTCACCTAAAGAGATCTCCATGGACGCGTGACGTCG | |

**Table S3 Butyl acetate production in the mutant strains with prophage deleted from the chromosome***

| | FJ-1201 48 h | FJ-1201 72 h | FJ-1301 48 h | FJ-1301 72 h |
|---|---|---|---|---|
| **EA** | <0.01 | <0.01 | <0.01 | <0.01 |
| **BA** | 19.3±0.5 | 19.7±1.7 | 19.1±0.5 | 19.4±0.7 |
| **BB** | 0.9±0.1 | 0.7±0.1 | 0.8±0.0 | 0.7±0.0 |
| **BA (aqueous phase)** | 0.6±0.1 | 0.6±0.1 | 0.5±0.0 | 0.5±0.1 |

*EA: ethyl acetate; BA: butyl acetate; BB: butyl butyrate. All values are in g/L.

**Table S4 FJ-1201 ester production using biomass hydrolysates as the substrate with or without supplementation of exogenous organic nitrogen source***

| | 0Y+0T# | 1Y+3T | 2Y+6T |
|---|---|---|---|
| **EA** | <0.01 | <0.01 | <0.01 |
| **BA** | 17.5±0.2 | 16.9±0.5 | 16.2±0.7 |
| **BB** | 0.1±0.0 | 0.2±0.1 | 0.2±0.1 |
| **BA (aqueous phase)** | 0.3±0.0 | 0.2±0.0 | 0.2±0.0 |

*EA: ethyl acetate; BA: butyl acetate; BB: butyl butyrate. All values are in g/L.
#Y=Yeast, T=Typtone; 0Y+0T: 0 g/L Y and 0 g/L T; 1Y+3T: 1 g/L Y and 3 g/L T; 2Y+6T: 2 g/L Y and 6 g/L T.

**Table S5. Project total capital investment ($ million) for the processes**

| | **Purchased Cost** | **Installed cost** |
|---|---|---|
| Feedstock handling | 15.5 | 26.4 |
| Pretreatment & hydrolysis | 18.0 | 29.8 |
| Fermentation | 36.6 | 55.2 |
| Production recovery & upgrading | 6.6 | 12.6 |
| Product & chemical Storage | 1.3 | 2.2 |
| Wastewater treatment | 59.8 | 59.8 |
| Electricity & steam generation | 34.4 | 68.4 |
| Utilities | 4.5 | 8.3 |
| **Total Installed Equipment cost** | | **262.7** |
| Other Direct Cost | | 17.1 |
| Total Direct Cost (TDC) | | 279.8 |
| Total Indirect Costs (TIC) | | 167.9 |
| **Fixed Capital Investment (FCI)** | | **447.6** |
| Land | | 1.8 |
| Working Capital | | 22.4 |
| **Total Capital Investment (TCI)** | | **471.8** |

**Table S6. Key operation parameters for the corn stover conversion and butyl acetate fermentation considered in the techno-economic analysis (TEA)**

| | |
|---|---|
| **Pretreatment and hydrolysis** | |
| Sodium hydroxide loading | 40 kg/MT dry corn stover |
| Deacetylation incubation temperature | 80 °C |
| Disk milling energy consumption | 212 kWh/MT dry corn stover |
| Enzymatic hydrolysis solids loading | 20% |
| Enzyme loading | 19 mg protein/g cellulose |
| Cellulose hydrolysis efficiency | 82% |
| Hemicellulose hydrolysis efficiency | 74% |
| **Fermentation** | |
| Hexadecane load | 1:1 by volume |
| Fermentation time | 96 hours |
| BA yield | 0.25 g BA/g consumed sugar |
| Butanol yield | 0.03 g butanol/g consumed sugar |
| Isopropanol yield | 0.04 g isopropanol/g consumed sugar |
| Additional nutrients | None (based on verified experiments) |

**Table S7. Summary of key raw material costs**

| Item | Cost ($) |
|---|---|
| **Raw materials and utilities** | |
| Corn stover (20% moisture) | 51.5/MT [a] |
| Sodium hydroxide | 116/MT [b] |
| Cellulase | 4,240/MT [a] |
| Hexadecane | 4,000/MT [b] |
| Wastewater treatment chemicals | 4,51/MT [b] |
| Boiler Chemicals | 8,336/MT [a] |
| Cooling tower chemicals | 3,668/MT [a] |
| Sulfuric acid | 41.0/MT [b] |
| Freshwater | 0.22/MT [a] |
| Electricity | 0.065/kWh [a] |
| Natural gas | 185/MT [b] |
| **Co-product credits** | |
| Butanol | 900/MT [b] |
| Isopropanol | 1150/MT [b] |
| Surplus Electricity | 0.065/kWh [b] |
| **Fixed Operating Costs** | |
| Labor costs | 2,500,000 [c] |
| Labor burden | 90% of labor cost |
| Maintenance | 3% of ISBL |
| Property insurance | 0.7% of fixed capital investment |

[a]The price of corn stover and other chemicals were from the previous literature including Humbird et al. (2011)[10], Chen et al. (2015)[9], and Dalle Ave and Adams (2018)[29];
[b]Data from different sources, including the ICIS chemical price report and industrial quotes;
[c]Assuming 50 employees with an average annual salary of $50,000 per employee.

**Table S8 Butyl acetate production in the engineered strains for the evaluation of the effect of gene codon optimization***

| | FJ-004 | FJ-007 |
|---|---|---|
| **EA** | <0.01 | <0.01 |
| **BA** | 5.5±0.5 | 5.0±0.2 |
| **BB** | 0.01±0.00 | < 0.01 |
| **Glucose consumption** | 79.3±0.1 | 76.7±2.4 |
| **Lactate** | 0.00 | 0.7±0.1 |
| **Acetate** | 0.3±0.00 | 0.2±0.1 |
| **Ethanol** | 1.0±0.0 | 1.0±0.0 |
| **Acetone** | 5.0±0.2 | 3.6±0.2 |
| **Butyrate** | 0.1±0.0 | 0.00 |
| **Butanol** | 7.8±0.0 | 9.5±0.7 |

* EA: ethyl acetate; BA: butyl acetate; BB: butyl butyrate. All values are in g/L.

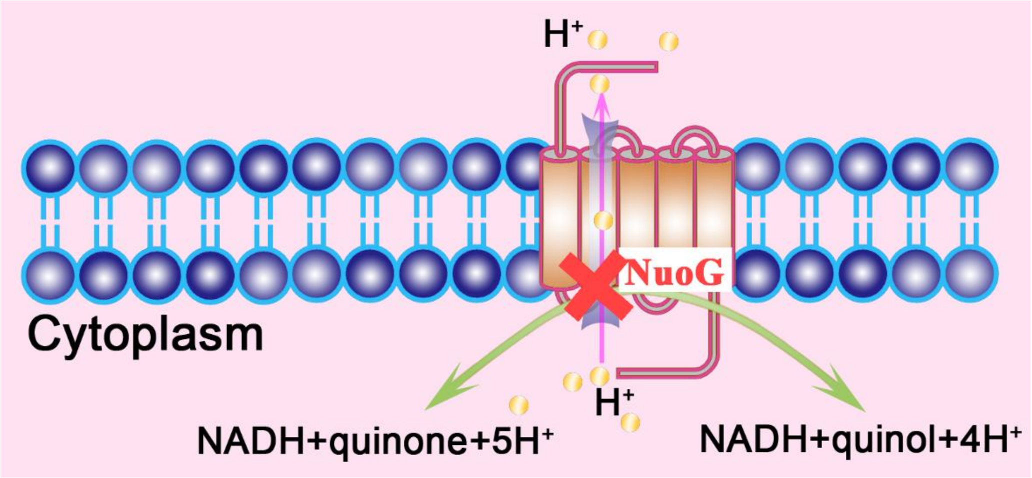


**Fig. S1 Schematic representation of the function of NuoG (quinone oxidoreductase subunit G) in clostridia**

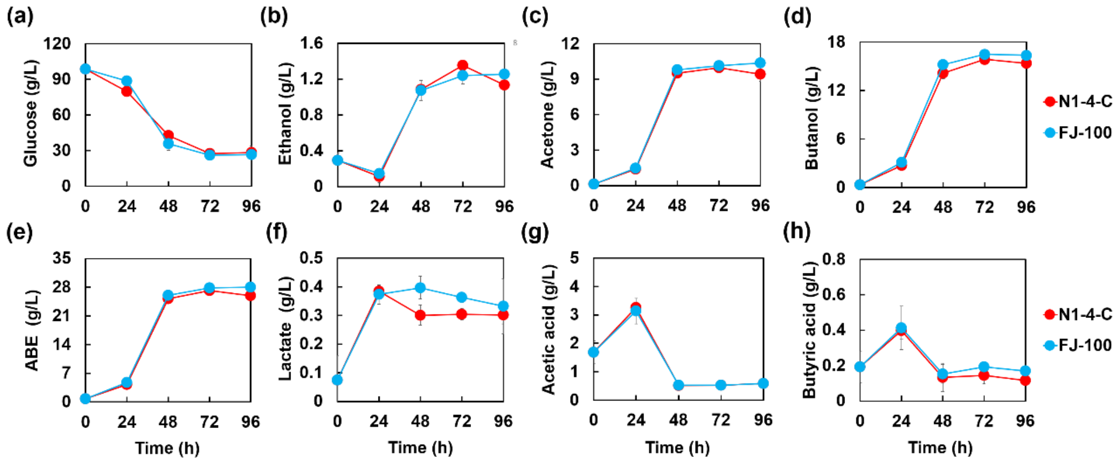


**Fig. S2 Fermentation results in serum bottles with *Clostridium saccharoperbutylacetonicum* N1-4-C and the mutant strains.** (a-h): ABE fermentation results for FJ-100 as compared to the control N1-4-C. The reported value is mean ± SD.

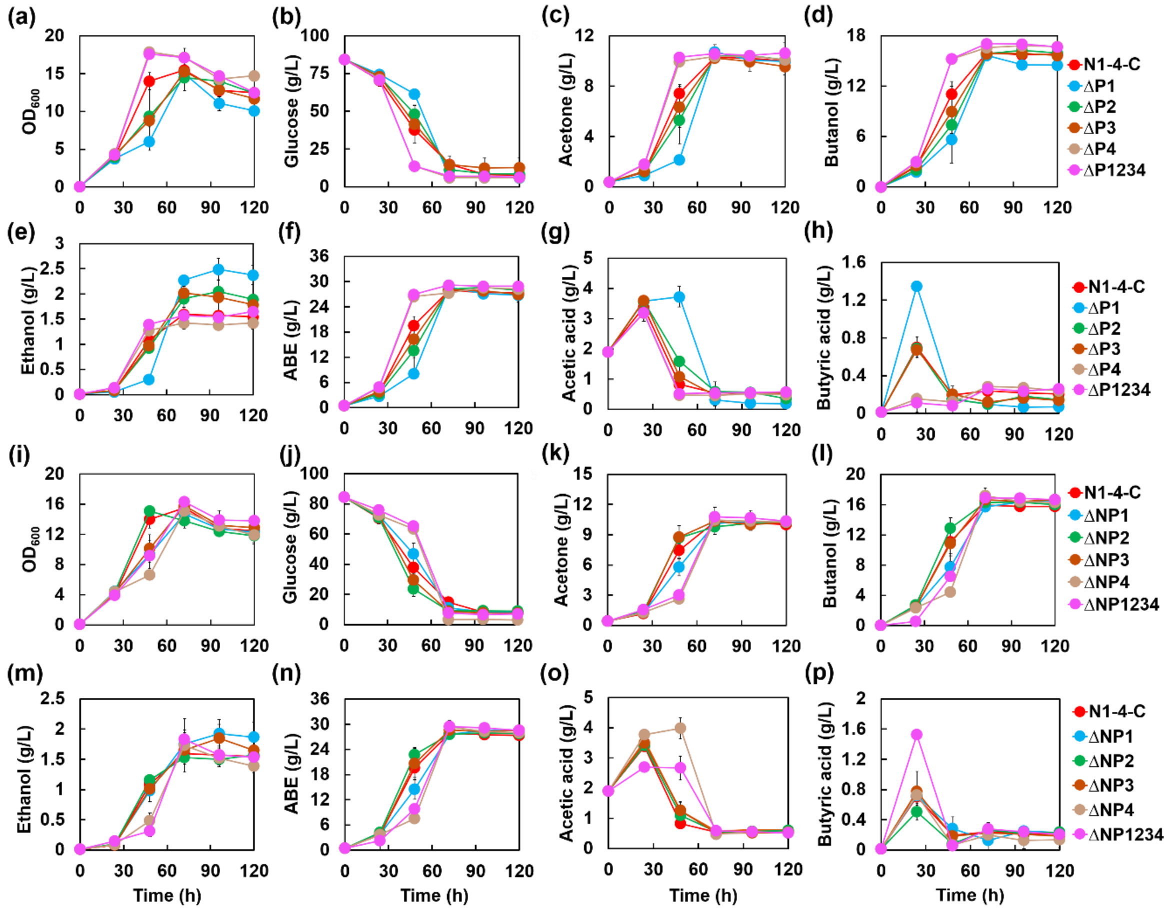


**Fig. S3 Fermentation results in serum bottles with *Clostridium saccharoperbutylacetonicum* N1-4-C and the mutant strains with prophage deleted.** (a-h): ABE fermentation results for ΔP1, ΔP2, ΔP3, ΔP4, ΔP1234 as compared to the control N1-4-C. (i-p): ABE fermentation results for ΔNP1, ΔNP2, ΔNP3, ΔNP4, ΔNP1234 as compared to the control N1-4-C. The reported value is mean ± SD.

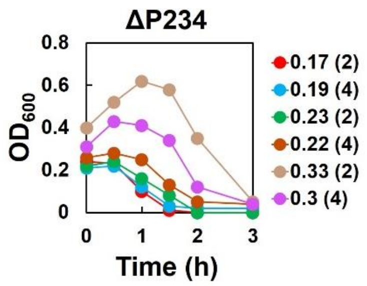


| Age of culture ($OD_{600}$) | Mitomycin C (μg/ml) | |
|---|---|---|
| | 2 | 4 |
| 0.1-0.2 | ++ | ++ |
| 0.2-0.3 | ++ | ++ |
| 0.3-0.4 | ++ | ++ |

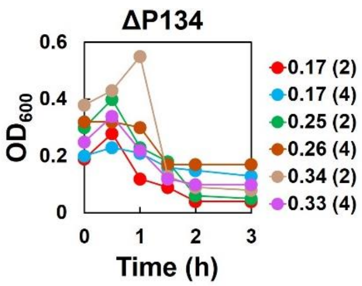


| Age of culture ($OD_{600}$) | Mitomycin C (μg/ml) | |
|---|---|---|
| | 2 | 4 |
| 0.1-0.2 | ++ | I |
| 0.2-0.3 | ++ | + |
| 0.3-0.4 | ++ | + |

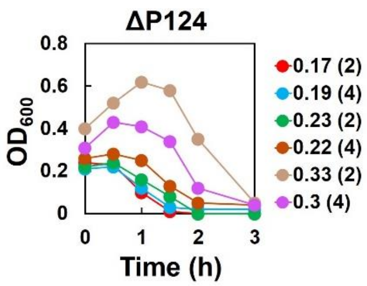


| Age of culture ($OD_{600}$) | Mitomycin C (μg/ml) | |
|---|---|---|
| | 2 | 4 |
| 0.1-0.2 | ++ | ++ |
| 0.2-0.3 | ++ | ++ |
| 0.3-0.4 | ++ | ++ |

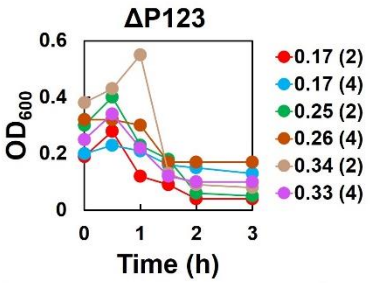


| Age of culture ($OD_{600}$) | Mitomycin C (μg/ml) | |
|---|---|---|
| | 2 | 4 |
| 0.1-0.2 | ++ | I |
| 0.2-0.3 | ++ | + |
| 0.3-0.4 | ++ | + |

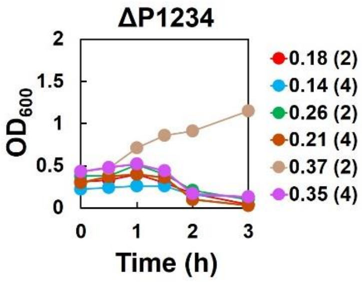


| Age of culture ($OD_{600}$) | Mitomycin C (μg/ml) | |
|---|---|---|
| | 2 | 4 |
| 0.1-0.2 | + | + |
| 0.2-0.3 | + | + |
| 0.3-0.4 | - | + |

**Fig. S4 Cell lysis in various prophage deletion mutants upon induction with different concentrations of mitomycin C.** "I" indicates growth inhibition; "-" indicates that there was no cell lysis; "+" indicates that most of the cells were lysed; "++" indicates that all the cells were lysed.

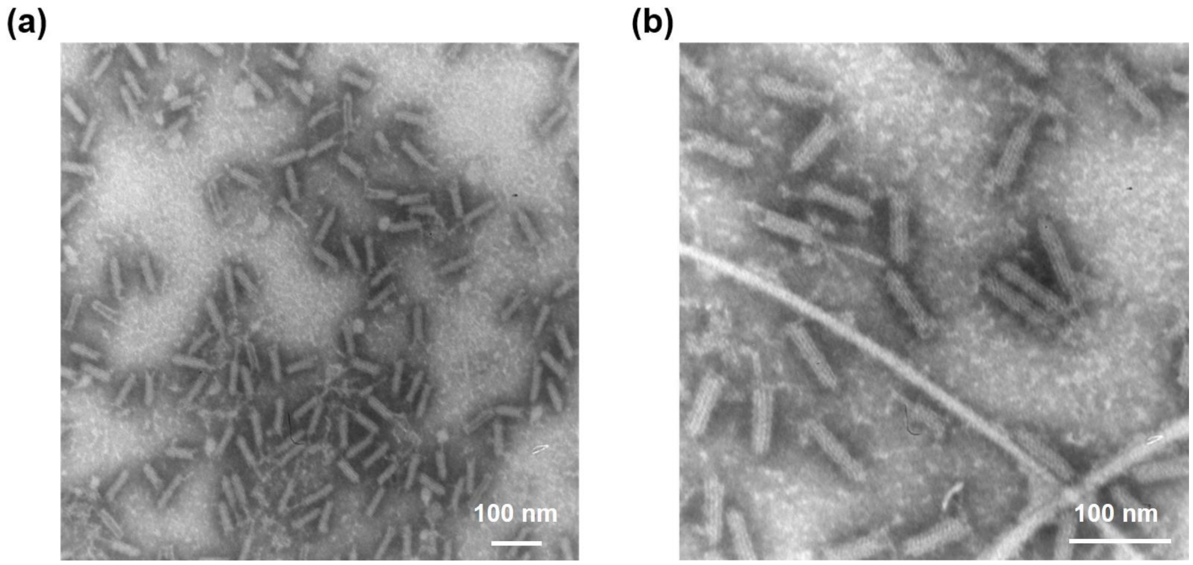


**Fig. S5 TEM picture of clostocin O.** The magnification of (a) was 100x, while the magnification of (b) was 200x.

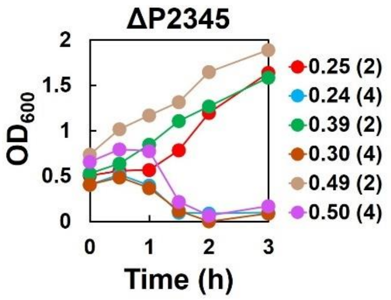


| Age of culture ($OD_{600}$) | Mitomycin C (μg/ml) | |
|---|---|---|
| | 2 | 4 |
| 0.2-0.3 | - | + |
| 0.3-0.4 | - | + |
| 0.4-0.5 | - | + |

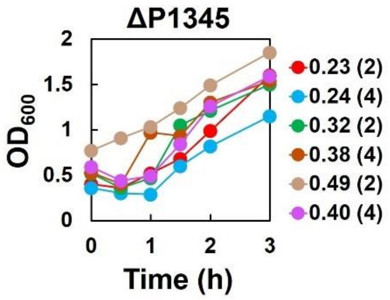


| Age of culture ($OD_{600}$) | Mitomycin C (μg/ml) | |
|---|---|---|
| | 2 | 4 |
| 0.2-0.3 | - | - |
| 0.3-0.4 | - | - |
| 0.4-0.5 | - | - |

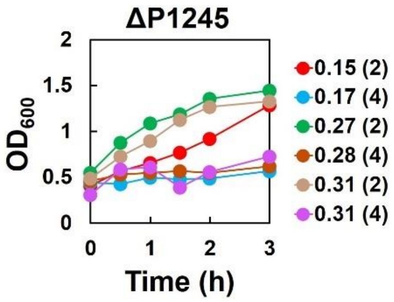


| Age of culture ($OD_{600}$) | Mitomycin C (μg/ml) | |
|---|---|---|
| | 2 | 4 |
| 0.1-0.2 | - | - |
| 0.2-0.3 | - | - |
| 0.3-0.4 | - | - |

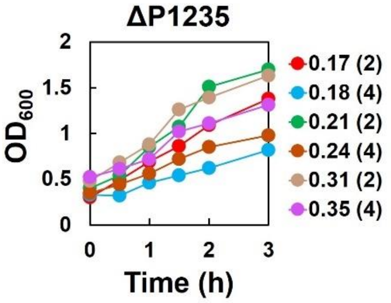


| Age of culture ($OD_{600}$) | Mitomycin C (μg/ml) | |
|---|---|---|
| | 2 | 4 |
| 0.1-0.2 | - | - |
| 0.2-0.3 | - | - |
| 0.3-0.4 | - | - |

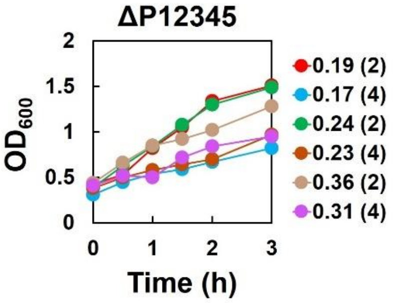


| Age of culture ($OD_{600}$) | Mitomycin C (μg/ml) | |
|---|---|---|
| | 2 | 4 |
| 0.1-0.2 | - | - |
| 0.2-0.3 | - | - |
| 0.3-0.4 | - | - |

**Fig. S6 Cell lysis in various prophage deletion mutants upon induction with different concentrations of mitomycin C.** "-" indicates that there was no cell lysis; "+" indicates that most of the cells were lysed.

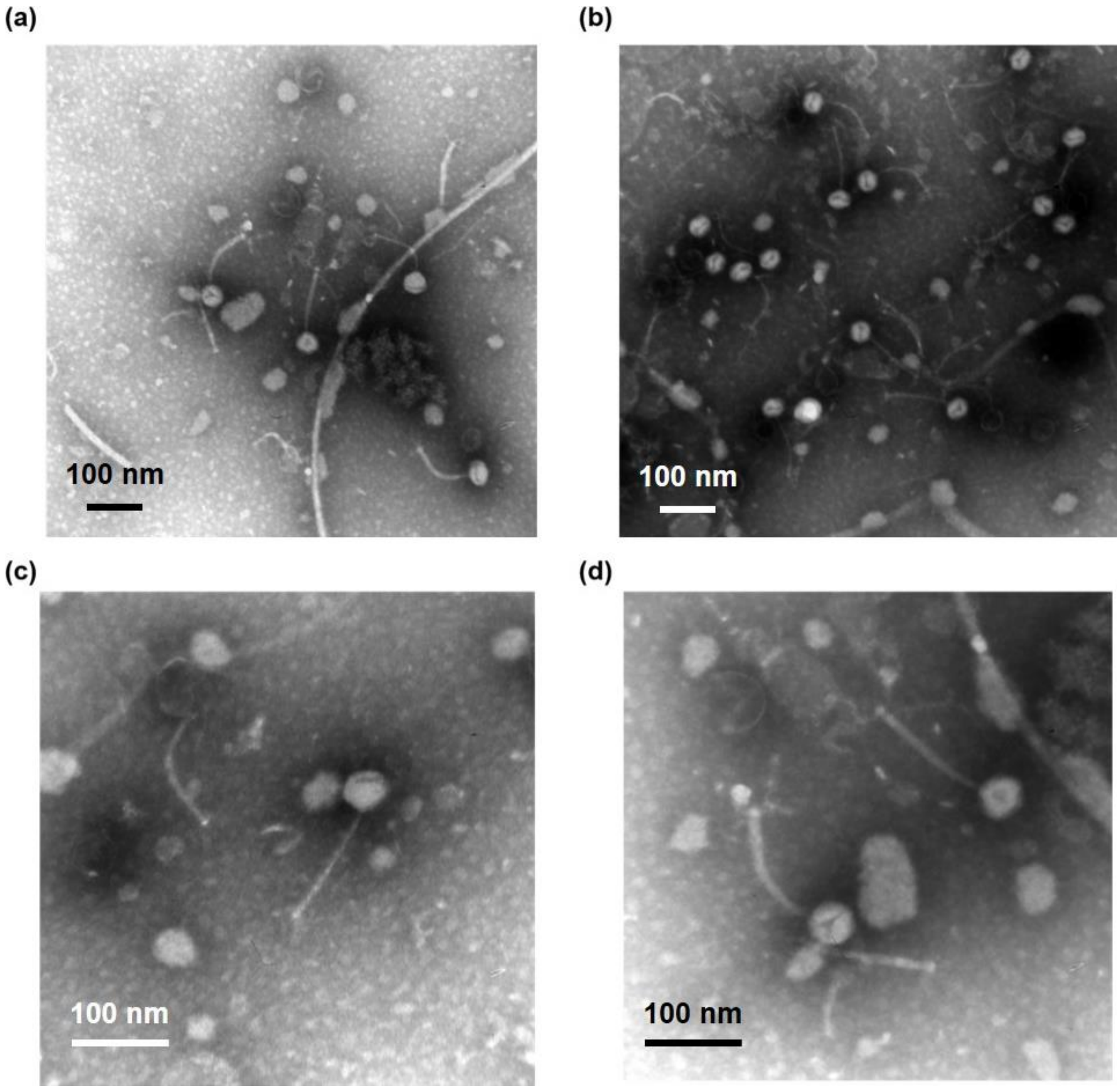


**Fig. S7 HM T phage observed in the ΔP2345 strain upon induction.** The magnification of (a, b) was 100x, while the magnification of (c, d) was 200x.

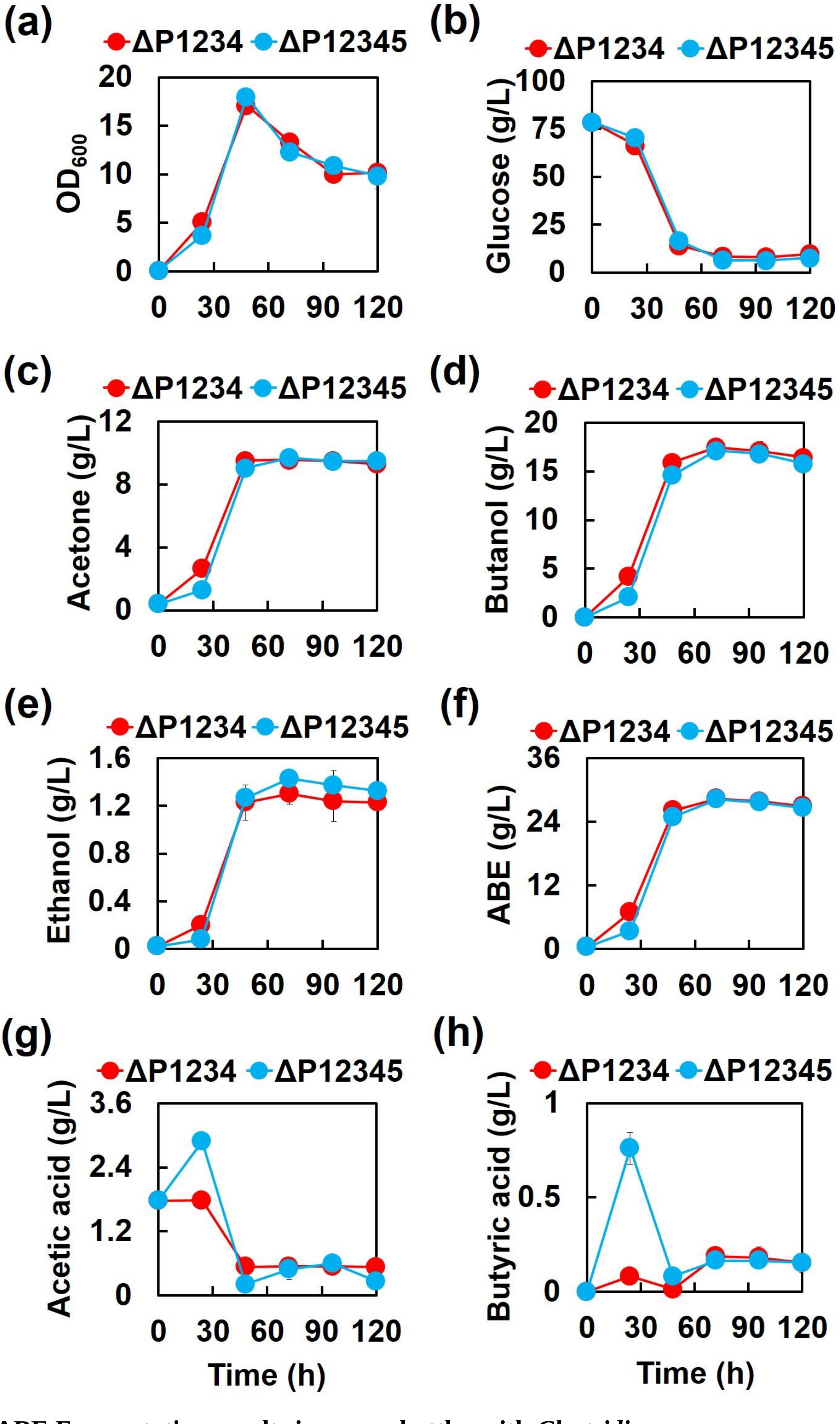


**Fig S8 ABE Fermentation results in serum bottles with *Clostridium saccharoperbutylacetonicum* ΔP1234 and ΔP12345.** The reported value is mean ± SD.

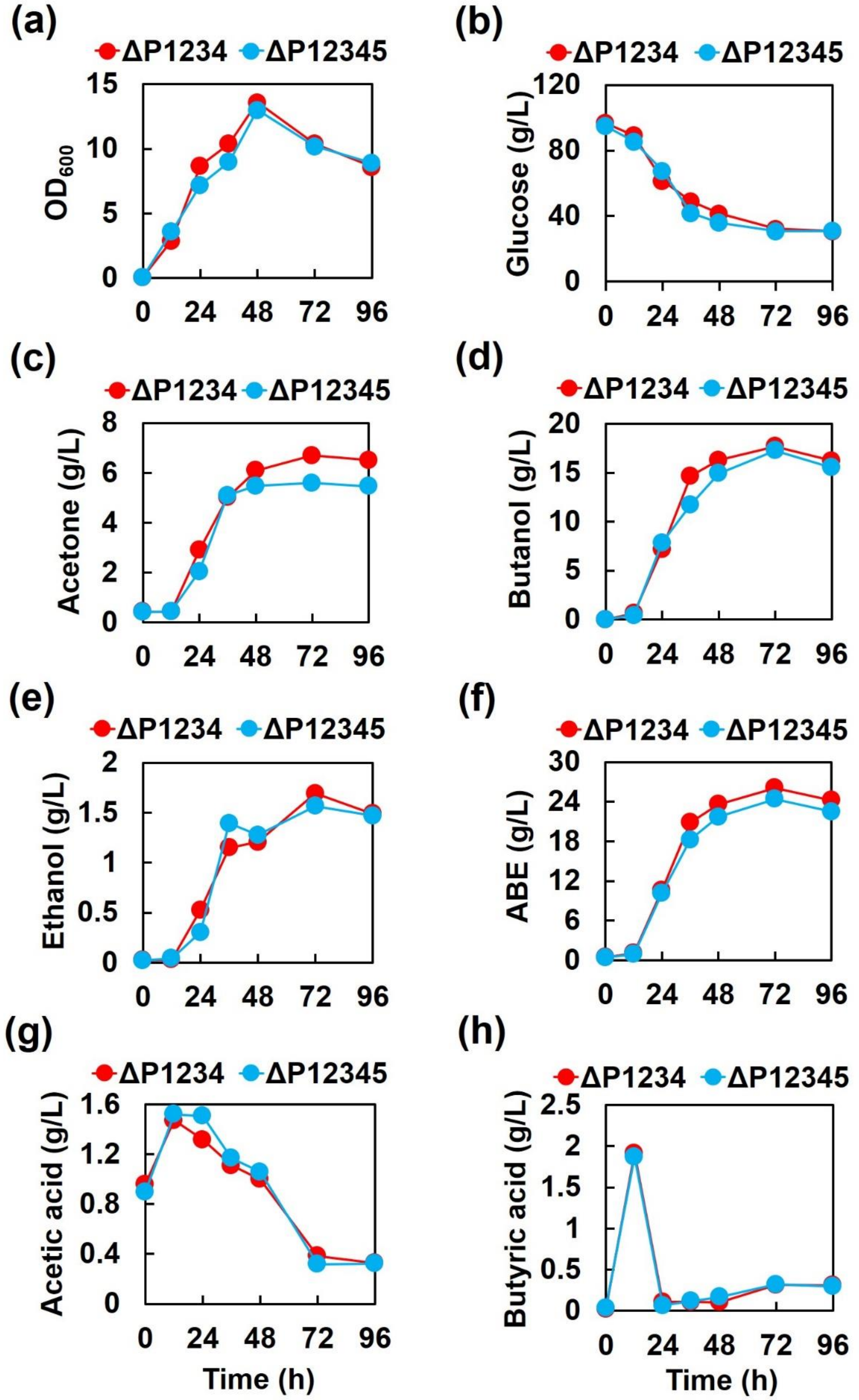


**Fig S9 ABE fermentation results in 500-mL bioreactors with *Clostridium saccharoperbutylacetonicum* ΔP1234 and ΔP12345.** The reported value is mean ± SD.

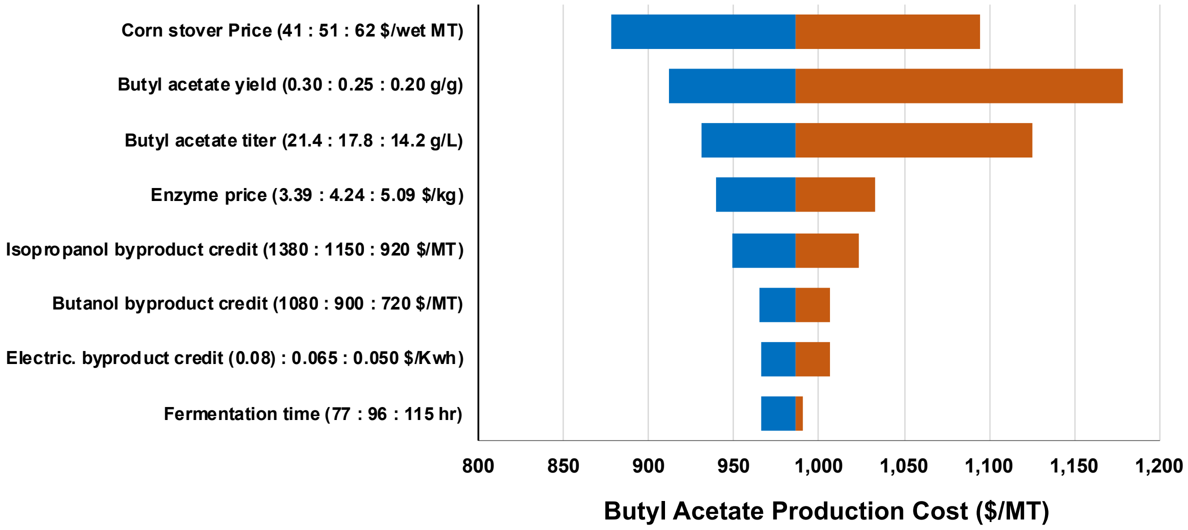


**Fig S10 Sensitivity of butyl acetate production cost to different parameters.** The numbers in brackets in Y-axis are the potential low, base and high values of each parameter.
